# Decoding molecular programs that define macrophage responses to tumor-derived cues

**DOI:** 10.64898/2026.06.05.730376

**Authors:** Katharina Hast-Sribike, Lukas Jonathan Häuser, Ariadna Acedo-Terrades, Rafael Riudavets-Puig, Julia Hau, Tiberiu Totu, Jonas Bossart, Angelica Patterson, Ekaterina Krymova, Vanesa Ayala-Nunez, Markus Rottmar, Katharina Maniura-Weber, Sonia Tugues, Marian Neidert, Bettina Sobottka, Marija Buljan

## Abstract

Tumor-associated macrophages (TAMs) comprise functionally diverse states that can suppress anti-tumor immunity and promote tumor progression, yet the tumor microenvironmental cues and signaling programs that generate these states remain incompletely defined. Here, we systematically stimulate primary human monocyte-derived macrophages with a panel of cytokines and metabolites abundant in the tumor microenvironment (TME), and profile their transcriptomic and phosphoproteomic responses to resolve stimulus-specific molecular programs. We observe that potassium (K^+^) and adenosine (Ado) stimulation, which accumulate in necrotic tumor cores, downregulate antigen-presentation genes and their master regulator *CIITA*. K^+^ stimulation results in the upregulated fibronectin 1 expression, associated with immunosuppressive, metastasis-promoting TAM subsets. Ado induces upregulated expression of tryptophan (Trp) catabolism genes, myeloid checkpoints and metallothioneins (MTs). Although MT-high TAM states have been recurrently observed across tumor single cell RNA sequencing studies, their function remains poorly defined. We show that elevated MT expression in tumor tissue is associated with shorter overall survival. By aligning *in vitro* transcriptomes with single-cell RNA sequencing (scRNA-seq) signatures from a pan-cancer TAM atlas, we identify significant similarities between several *in vitro* states and clinically observed TAM populations, with Ado-stimulated macrophages closely resembling a MT-expressing TAM cluster. Overall, this work provides a systematic molecular context linking tumor microenvironmental cues to clinically relevant TAM states and offers a framework for recapitulating their functions *in vitro*.

**STATEMENT OF SIGNIFICANCE:** This study explores how cytokines and metabolites from the tumor microenvironment shape macrophage molecular phenotypes and lead to the upregulation of clinically relevant marker genes and recapitulation of functional states of interest.

## INTRODUCTION

TAMs play a central role in cancer progression, exerting both immunosuppressive and tumor-promoting functions. TAM-mediated immunosuppression is enabled through the secretion of cytokines [1-3], loss of antigen presenting functions [4], expression of myeloid checkpoints and consequent inhibition of macrophage-mediated phagocytosis of malignant cells [5, 6] as well as by depriving T cells of essential metabolites such as Trp [7, 8]. In parallel, tumor growth is promoted through TAM-induced angiogenesis and extracellular matrix (ECM) organization [9, 10], direct interactions with cancer stem cells [11, 12], as well as the facilitation of metastatic niche formation [13, 14]. In contrast to these functions, distinct TAM subpopulations play a critical role in anti-tumor immunity through presentation of tumor antigens, tumor cell phagocytosis and immune cell recruitment [15-17]. This functional heterogeneity and remarkable plasticity, along with their innate tumor-homing capacity, position TAMs as attractive targets for next-generation immunotherapies [18-20].

scRNA-seq studies of patient tumor resections have revealed a diverse spectrum of clinically relevant TAM subpopulations [21-23]. Among others, macrophages expressing the well-established immunosuppressive markers CD206 (encoded by *MRC1*) and CD163 associate with tumor progression [24, 25]. Beyond these canonical polarization markers, recent studies have identified transcriptional markers and molecular programs that distinguish macrophage states in patient the TME. Lipid-associated TREM2^+^ TAMs and hypoxic SPP1^+^ TAMs are among the most recurrent immunosuppressive subtypes found in solid tumors, whereas CXCL9^+^ TAMs represent a common inflammatory subtype [15, 20, 26-29]. Additionally, clinically relevant TAM states can be identified by the expression surface receptors, metabolic genes, or structural proteins. MARCO^+^ TAMs constitute an immunosuppressive subset characterized by high scavenger activity [30, 31], while upregulation of *APOE* or *APOC1* genes is linked to metabolic rewiring favoring lipid metabolism, immunosuppressive activation, and resistance to immune checkpoint blockade [32-35]. FN1^+^ TAMs have recently emerged as a protumoral state promoting metastatic progression and recurrence [36, 37]. However, the clinical implications of these TAM states are highly context-dependent and vary across tumor types, disease stages, and therapeutic settings [20]. For instance, TREM2^+^ TAMs have been associated with unfavorable prognosis in several cancers [28, 38], but also with favorable outcome or therapy response in others [39]. Although preclinical mouse studies suggest that targeting TREM2 can reduce tumor growth and enhance antitumor immunity [38], early clinical evaluation of anti-TREM2 therapy showed only limited efficacy [40]. These findings underscore both the translational limitations of extrapolating from mouse models to human TAM biology and the need to move beyond marker-defined TAM categories. Moreover, although scRNA-seq studies have revealed clinically relevant TAM heterogeneity, the local TME stimuli that drive the polarization of these states and the effector programs they acquire remain insufficiently understood.

The TME provides a highly dynamic niche in which TAM polarization is shaped by concentration and combinations of received stimuli, as well as exposure timing and duration [41, 42]. Consequently, *in vitro* macrophages do not fully recapitulate clinically relevant TAM states, but remain essential for dissecting stimulus-response relationships, mapping physiologically and pathologically relevant signaling routes, prioritizing markers of interest and supporting preclinical drug development [43, 44]. Clinically relevant TAM phenotypes likely arise from combinatorial activation of a finite set of signaling pathways. Importantly, even though the knowledge on signaling pathways is commonly extrapolated from one system to another, signaling routes are cell-type specific [45, 46], rendering their definition in primary human macrophages critical. TME-derived metabolites and cytokines are key regulators of macrophage polarization and function [47]. Hypoxia and ne-crosis elevate extracellular Ado and K^+^ concentrations, both of which were reported to promote immunosuppressive TAM programs [48-52]. Ado signaling through the TAM A2A receptor (A2AR) impairs T- and NK-cell immunity, whereas A2AR blockade reverses TAM-mediated immunosuppression and enhances immunotherapy responses [53]. Similarly, inhibition of the K^+^ channel Kir2.1 repolarizes TAMs toward an antitumor phenotype in mice [48]. In addition to metabolites, TME-abundant cytokines, such as Transforming Growth Factor β (TGFβ), skew macrophages toward tumor-promoting phenotypes [54]. TGFβ receptor inhibition can induce pro-inflammatory shifts in TAMs and improve therapeutic responses [55-57]. Although these studies highlight the therapeutic potential of targeting TME-derived stimuli and their receptors, the underlying signaling networks in primary human macrophages remain incompletely defined and clinical translation of TAM-modulating drugs remains limited [20, 58]. A more detailed charting of these networks may be crucial for expanding the repertoire of druggable targets and guiding rational design of TAM-specific therapeutic strategies.

Recent studies have taken a major leap in the systematic mapping of immune-cell responses to single stimuli: The Immune Dictionary profiled cytokine-induced immune response in mouse lymph nodes using scRNA-seq [59], while the Human Cytokine Dictionary analogously mapped cytokine-induced transcriptional responses of primary human peripheral blood mononuclear cells [60]. However, comparable systematic profiling of human primary monocyte-derived macrophages remains limited, with previous *in vitro* studies assessing only a small number of stimuli using microarray-based approaches [61].

In this study, we systematically stimulate primary human macrophages with tumor-abundant soluble factors and profile their multi-omics signature to generate macrophage states that replicate specific TAM programs. These analyses identify stimulus-induced shifts in inflammatory, metabolic, antigen-presentation, and ECM remodeling programs, including TGFβ-driven lipid metabolism signatures and suppression of inflammation, K^+^-induced skewing toward interferon (IFN) responses and *FN1* expression. We further observe that both Ado and K^+^ stimulation cause a downregulation of antigen presentation Major Histocompatibility Class-II (MHC-II) genes as well as their regulator *CIITA*, potentially limiting their antigen presentation capacity. Additionally, Ado stimulation results in the upregulated expression of myeloid checkpoint, Trp catabolism and MT genes. To contextualize these *in vitro* generated macrophage sets, we align their transcriptomes to a pancancer scRNA-seq atlas of patient TAM signatures [21]. Several *in vitro* generated states share significant phenotypic similarities with TAM populations observed in patients. AdoMacs share features of a MT-expressing TAM clusters observed in patients, secrete elevated levels of immunosuppressive cytokines and display reduced susceptibility to the macrophage reprogramming drug Resiquimod (R848) compared to canonical M2-like macrophages. Collectively, this study provides a multi-omics resource linking tumor microenvironmental cues to clinically relevant TAM states and offers an experimental framework for the rational design of precision therapies targeting specific TAM states.

## RESULTS

### Generation of a spectrum of *in vitro* macrophage states

We curated a panel of cytokines and metabolites previously reported to be (i) abundant in the TME and (ii) able to initiate macrophage polarization (see Methods, Table 2). To reflect the diversity of macrophage-activating cues within the TME, the stimulus panel included metabolic, ionic, and hypoxia-associated stress signals such as γ-aminobutyric acid (GABA) [1, 62], Ado [49, 51-53], lactate (Lac) [2, 63], succinate [64, 65], K^+^ [48], and oleate [66, 67], immunomodulatory cytokines and chemokines such as leukemia inhibitory factor (LIF) [68-70], oncostatin M (OSM) [71], TGFβ [72, 73], interleukin (IL)-33 [74, 75], C-C motif chemokine ligand 5 (CCL5) [76], and CCL2 [77], as well as the vascular and stromal remodeling factors vascular endothelial growth factor (VEGF) [78] and bone morphogenetic protein 7 (BMP7) [79]. Based on reported synergistic effects, we also included a combined TGFβ + IL-33 condition [80]. To assess whether signaling routes induced by individual stimuli are recapitulated in macrophages exposed to a complex tumor-derived soluble factor milieu, we additionally generated tumor-conditioned media (TCMs) from four fresh resections of central nervous system neoplasms (see Methods, Table 1) by dissecting tumor tissues into organoids and culturing them in macrophage medium for three days. Then, we differentiated buffy coat–derived human monocytes into macrophages using M-CSF and exposed them for 24 h to single or combined stimuli at literature-reported concentrations, or to TCMs at a 1:1 ratio, to generate macrophage states that potentially replicate specific stimulus-induced TAM programs. (**Fig. 1a**). Untreated macrophages served as a baseline control, while canonical *in vitro* polarization conditions were used as reference states for benchmarking the experimental approach: Inflammatory M1Macs stimulated with lipopolysaccharides (LPS) and IFNγ, M2aMacs stimulated with IL-4 and IL-13, and M2cMacs stimulated with IL-10.

**Table 1.** Patient overview of resected tumor tissue.

| Patient | Tumor type | Gender | Age |
| --- | --- | --- | --- |
| P1 | Meningioma | female | 65 |
| P2 | Meningioma | male | 60 |
| P3 | Vestibular Schwannoma | male | 72 |
| P4 | Meningioma | female | 43 |

**Table 2.** Macrophage polarization medium composition.

| Condition | Stimulus | Category | Concentration | Supplier | Ref |
| --- | --- | --- | --- | --- | --- |
| Untreated | - | Control | - |  |  |
| LPS + IFN-γ | Lipopolysaccharides from Salmonella enterica serotype enteritidis/<br>Human IFN-gamma Recombinant Protein (16.8 kDa) | Cytokine | LPS: 100 ng/mL<br>IFNγ: 20 ng/mL | Merck/<br>PeproTech | |
| IL-4 + IL-13 | Human IL-4 Recombinant Protein (15.1 kDa)/<br>Human IL-13 Recombinant Protein (12.6 kDa) | Cytokine | 20 ng/mL each | Gibco |  |
| IL-10 | Human IL-10, Animal-Free Recombinant Protein (18.6 kDa) | Cytokine | 20 ng/mL | PeproTech |  |
| LIF | Human Leukemia inhibitory factor Recombinant Protein (19.7 kDa) | Cytokine | 20 ng/mL | PeproTech | [69] |
| OSM | Recombinant Human Oncostatin M (25.7 kDa) | Cytokine | 20 ng/mL | PeproTech | [71] |
| TGFβ | Recombinant Human Transforming Growth Factor -β1 (25.0 kDa) | Cytokine | 20 ng/mL | PeproTech | [72] |
| VEGF | Human vascular endothelial growth factor 165 Recombinant Protein (38.2 kDa) | Cytokine | 20 ng/mL | PeproTech | [78] |
| IL-33 | Recombinant Human IL-33 Recombinant Protein (17.9 kDa) | Cytokine | 20 ng/mL | PeproTech | [74] |
| TGFβ/IL-33 | See above | Cytokine | 20 ng/mL each | PeproTech | [80] |
| CCL5 | Human CCL5 (RANTES) Recombinant Protein (7.8 kDa) | Cytokine | 20 ng/mL | PeproTech | [76] |
| CCL2 | Human CCL2 (MCP-1) Recombinant Protein (8.6 kDa) | Cytokine | 20 ng/mL | PeproTech | [77] |
| BMP-7 | Human bone morphogenetic protein 7 Recombinant Protein (26.4 kDa) | Cytokine | 20 ng/mL | PeproTech | [79] |
| GABA | γ-Aminobutyric acid (CAS-Nr. 56-12-2) | Metabolite | 1 mM | Sigma | [1] |
| Adenosine | Adenosine (CAS-Nr. 58-61-7) | Metabolite | 100 μM | Sigma | [52] |
| Lactate | DL-Lactic acid (CAS-Nr. 50-21-5) | Metabolite | 5 mM | Sigma | [63] |
| Succinate | Succinic acid (CAS-Nr. 110-15-6) | Metabolite | 1 mM | Sigma | [64] |
| K<sup>+</sup> | Potassium chloride (CAS-Nr. 7447-40-7) | Metabolite | 40 mM | Sigma | [48] |
| Oleate | Oleic acid (Cas-Nr. 112-80-1) | Metabolite | 100 μM | Sigma | [67] |
| TCM1 | Tumor-conditioned medium (P1) | Complex | 1:1 ratio |  |  |
| TCM2 | Tumor-conditioned medium (P2) | Complex | 1:1 ratio |  |  |
| TCM3 | Tumor-conditioned medium (P3) | Complex | 1:1 ratio |  |  |
| TCM4 | Tumor-conditioned medium (P4) | Complex | 1:1 ratio |  |  |

**Fig. 1:**
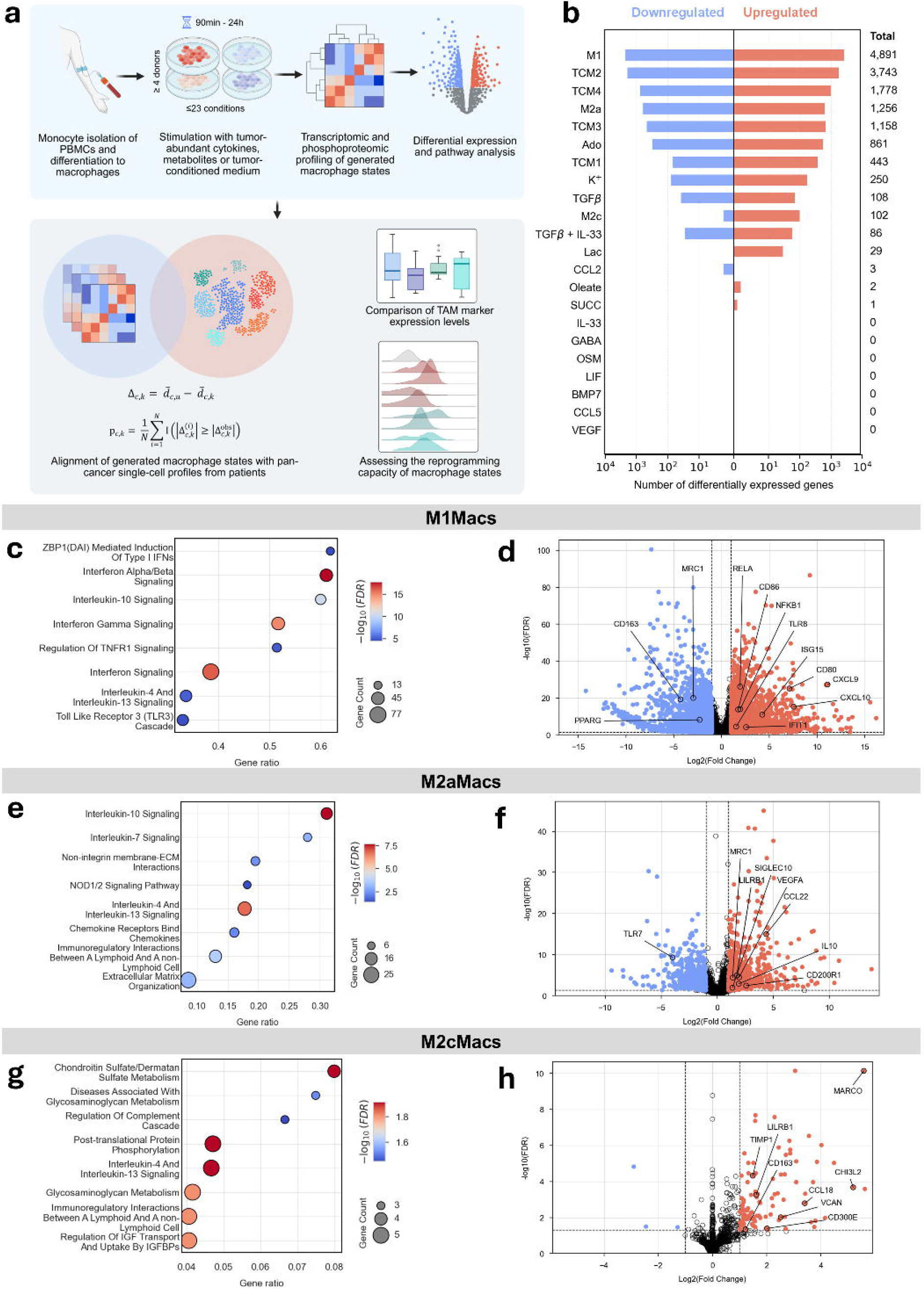
Generation of a spectrum of human primary *in vitro* macrophage states. **a**, Schematic overview of the experimental and computational workflow. First row: macrophage polarization, multi-omics characterization and analysis of the generated datasets for the representative macrophage *in vitro* states; second row, alignment of *in vitro* transcriptomes with single cell macrophage transcriptomes from patient tumors and assessment of the clinical relevance of generated macrophage states. **b**, Quantitative representation of the total number of significant DEGs across the conditions, 24 h after stimulation. The individual stimulus-induced transcriptional responses were compared to untreated controls and DEGs are defined when the transcripts levels have a log2FC ≥ |1| and an FDR ≤ 0.05. **c**, Dot plot depicting top over-represented Reactome pathways enriched among DEGs in macrophages stimulated with LPS and IFNγ (M1Macs) in comparison to untreated macrophages. Dot size indicates the number of DEGs per term; color scale reflects FDR; gene ratio□= □number of DEGs with a Reactome term/total number of genes annotated with the term. **d**, Volcano plot illustrating gene expression changes in M1Macs when compared to untreated macrophages. The X axis depicts log2FC of gene expression levels and the Y axis the corresponding −log_10_ (FDR) values. A log2FC ≥ |1| and an FDR ≤ 0.05 were used as thresholds for significant hits. DEGs are colored blue and red, depending on the directionality of the expression change, i.e. under- or overexpression compared to untreated macrophages. **e**, Dot plot depicting top over-represented Reactome pathways enriched among the DEGs in macrophages stimulated with IL-4 and IL-13 (M2aMacs). DEGs are defined as above. **f**, Volcano plot illustrating gene expression changes in M2aMacs when compared to untreated macrophages. Graph axes, colors and thresholds are as in d. **g**, Dot plot depicting top over-represented Reactome pathways enriched among the DEGs in macrophages stimulated with IL-10 (M2cMacs). DEGs are defined as above. **h**, Volcano plot illustrating gene expression changes in M2cMacs when compared to untreated macrophages. Graph axes, colors and thresholds are as in d.

Next, we performed bulk transcriptomic profiling of generated macrophage states from four healthy donors to map stimulus-induced transcriptional shifts. Donor effects were included as a covariate in the analyses (Suppl. Fig. 2). Each stimulated condition was compared to untreated macrophages when performing the differential expression analysis (Suppl. Table 1). Differentially expressed genes (DEGs) were defined by an expression level of log2FC ≥ |1| with an FDR ≤ 0.05. Among tested conditions, stimulation with LPS+IFNγ, IL-4+IL-13, IL-10, Ado, K^+^, TGFβ, Lac, oleate and TCMs elicited a pronounced transcriptional responses (Fig. 1b, Suppl. Fig. 1). For these conditions, we then performed overrepresentation analysis on DEGs using Reactome [81] and Gene Ontology [82] annotations, as well as gene set enrichment analysis (GSEA) on ranked gene lists using MSigDB Hallmark [83] annotations to identify molecular pathways and functional programs enriched in each macrophage state (Suppl. Table 2, 3).

**Fig. 2:**
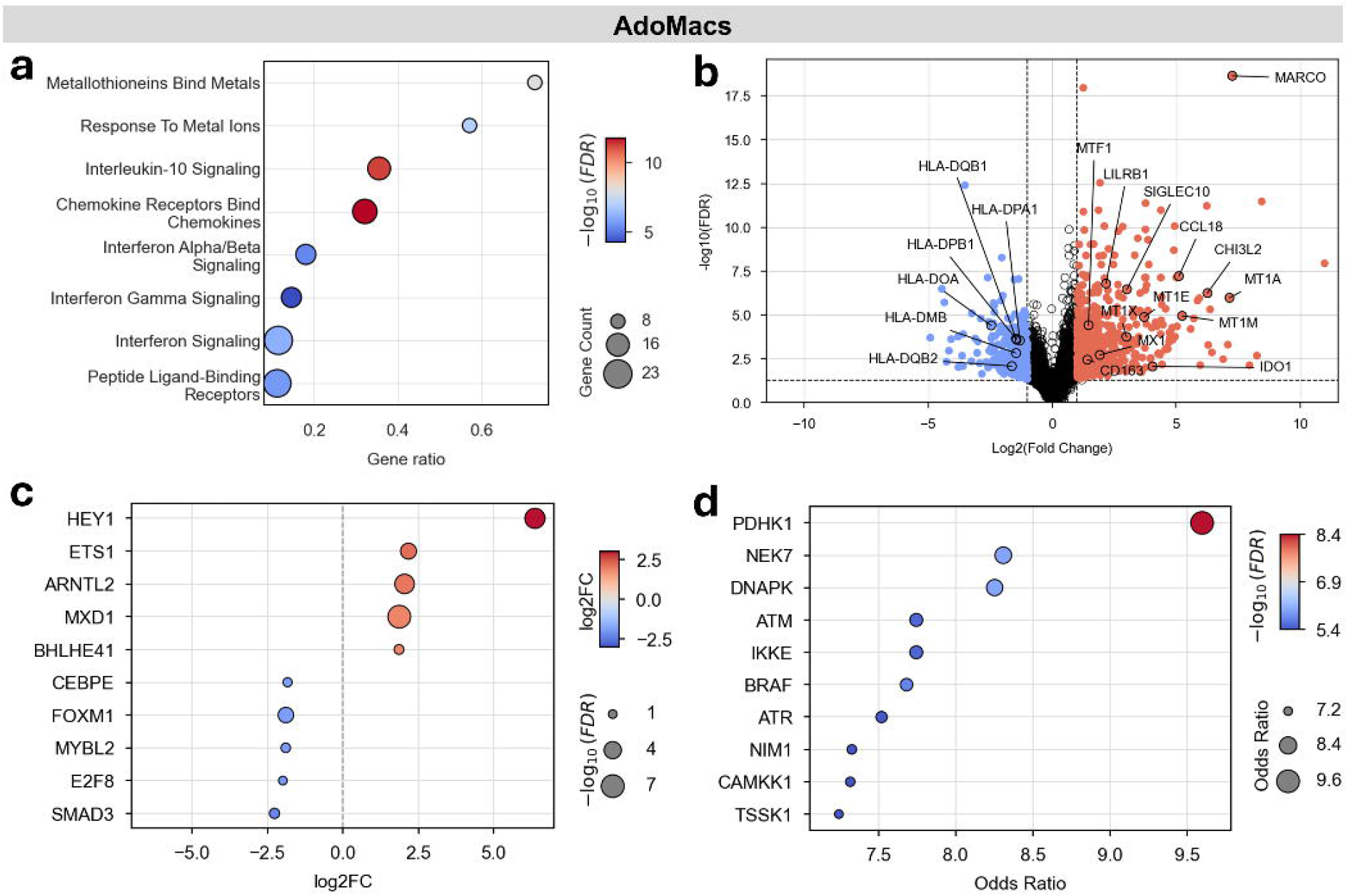
Characterization of the macrophage phenotypic response to high extracellular adenosine. **a**, Dot plot depicting top over-represented Reactome pathways among DEGs in macrophages stimulated with adenosine (AdoMacs). DEGs are defined in comparison to untreated macrophages with the same thresholds as detailed in Figure 1. **b**, Volcano plot illustrating gene expression changes in AdoMacs when compared to untreated macrophages. Graph axes, colors and thresholds are as in Figure 1. **c**, Dot plot illustrating top differentially up- and downregulated transcription factor genes in AdoMacs when the gene expression levels are compared to untreated macrophages. A less restricted threshold of log2FC ≥ |0.5| and an FDR ≤ 0.05 were used to indicate significant TFs. **d**, Dot plot depicting kinases that are inferred to be upstream regulators of the significantly differentially upregulated phosphopeptides in AdoMacs. The odds ratio indicates the enrichment of substrates associated with a given kinase among the input proteins relative to their expected occurrence in the background set.

All conditions exhibited reduced expression of proliferation-associated gene sets, consistent with an enhanced differentiation after the applied stimuli (Suppl. Table 2) [84, 85]. Macrophages stimulated with LPS and IFNγ (M1Macs) exhibited upregulation of its marker genes *CD80* and *CD86* as well as strong upregulation of pathways related to inflammation and chemotaxis, IFN- and Toll-like receptor (TLR)/NFκB-signaling (**Fig. 1c, d**, Suppl. Fig. 3a, Suppl. Notes) [42, 86]. Comparison of macrophages stimulated with IL-4 and IL-13 (M2aMacs) with untreated macrophages showed upregulation of the marker CD206 (encoded by *MRC1*) typical for the M2a-like state [86] as well as upregulation of genes associated with IL-4/ IL-13/IL-10 signaling pathways, extracellular matrix (ECM) remodeling, and inhibitory cell-cell interactions (**Fig. 1e, f**, Suppl. Notes) [63, 87]. Compared to untreated macrophages, macrophages stimulated with IL-10 (M2cMacs) showed upregulation of pathways associated with immunosuppressive IL-4/IL-13 cytokine signaling (**Fig. 1g**, Suppl. Notes), ECM remodeling through glycosaminoglycan metabolism, and interactions with lymphoid cells, consistent with literature on M2c-like macrophages [87, 88]. Among the most highly upregulated genes in M2cMacs were immunosuppressive TAM marker genes *MARCO* [30, 31] and *CD163* (**Fig. 1h**) Taken together, verification of the expected molecular phenotypes typical for *M1, M2a* and *M2c in vitro* states suggests that despite substantial donor interindividual variability, strong stimuli initiate robust molecular changes in primary human macrophages [60, 86, 89].

**Fig. 3:**
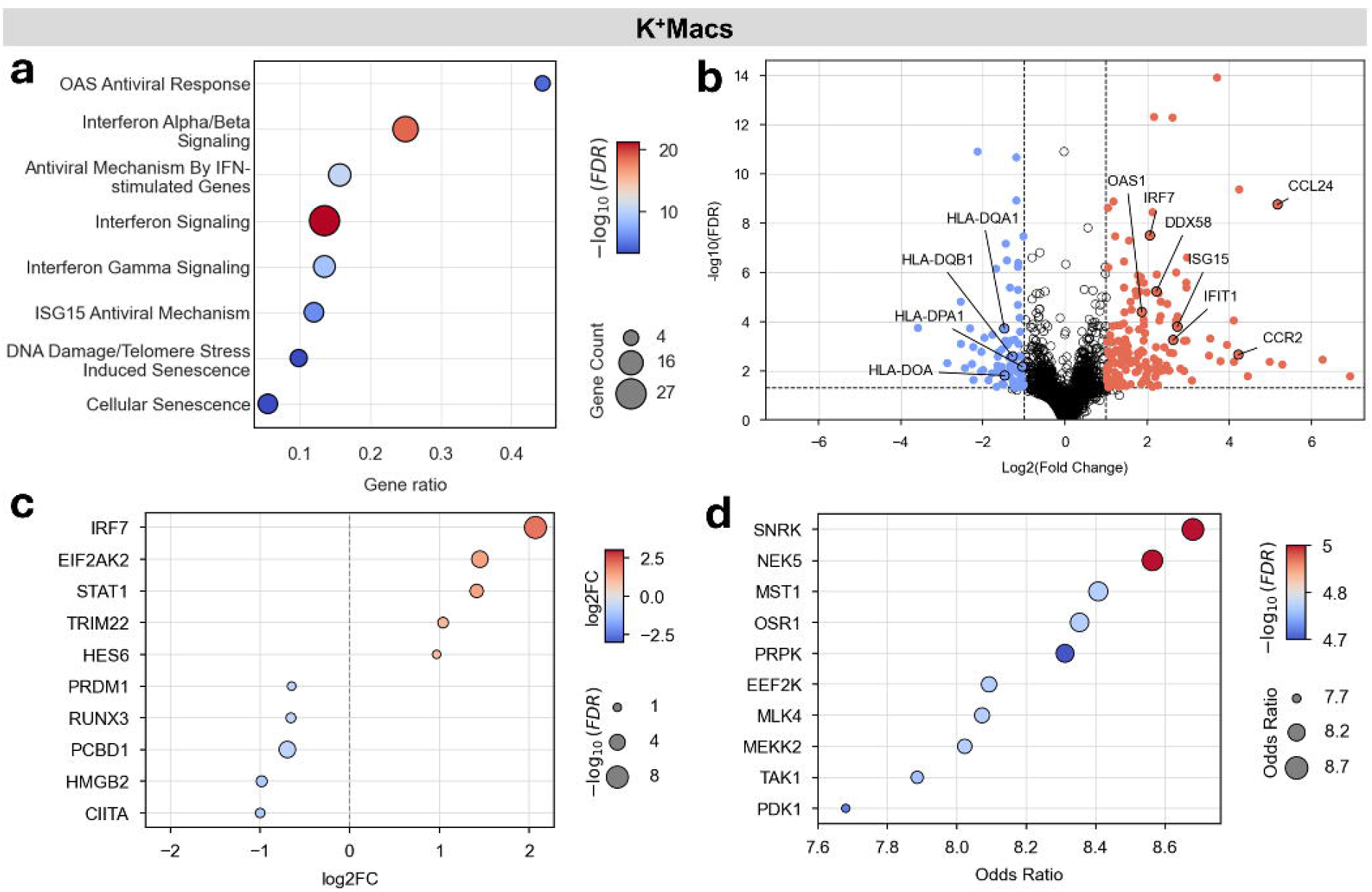
Characterization of the macrophage molecular responses to high extracellular K^+^. **a**, Dot plot depicting over-represented Reactome pathways among DEGs in macrophages stimulated with potassium chloride (K^+^Macs). DEGs are defined in comparison to untreated macrophages with the same thresholds as detailed in Figure 1. **b**, Volcano plot illustrating gene expression changes in K^+^Macs when compared to untreated macrophages. Graph axes, colors and thresholds are as in Figure 1. **c**, Dot plot illustrating top differentially up- and downregulated transcription factor genes in K^+^Macs when the gene expression levels are compared to untreated macrophages. A threshold of log2FC ≥ |0.5| and an FDR ≤ 0.05 were used to indicate significant TFs. **d**, Dot plot depicting kinases that are inferred to be upstream regulators of the significantly differentially upregulated phosphopeptides in K^+^Macs.

### Adenosine and K^+^ induce tolerogenic macrophage programs

Treatment with Ado induced notable changes in macrophage transcriptomes (**Fig. 1c**), most prominently, we upregulated cellular pathways associated with metabolism of metal ions (**Fig. 2a**). This trend was driven by the strong upregulation of genes in the MT family (*MT1A, MT1DP, MT1E, MT1F, MT1G, MT1H, MT1JP, MT1L, MT1M, MT1X, MT2A*) as well as the Metal-responsive transcription factor (*MTF1*, log2FC > 1.4, FDR < 0.001, Suppl. Table 4) (**Fig. 2b**). MT-expressing TAM states have been recurrently identified in scRNA-seq studies of patient tumors; however, their functional roles within the TME remain poorly characterized [21, 23, 90]. MTs are critical regulators of immune processes and act as zinc reservoirs [91, 92]. Their expression can be induced by hypoxia and they can promote cell survival in stress conditions by mitigating oxidative stress and scavenging reactive oxygen species [93-95]. Upregulated genes in adenosine-stimulated macrophages (AdoMacs) were also enriched in pathways related to chemotaxis and inflammation (**Fig. 2a**, Suppl. Fig. 3d), as illustrated by the expression of *MX1, MX2, IFIT1, IFIT2, IFIT3, CXCL10*, and *IL1B* (**Fig. 2b**, Suppl. Table 2). However, simultaneously, they upregulated the immunosuppressive cytokines *CCL18* and *CCL23* (Suppl. Table 1), as well as pathways related to STAT3 and IL-10 signaling (**Fig. 2a**) [3, 96]. This transcriptional pattern is reminiscent of macrophage states induced by hypoxia, which can involve the concurrent upregulation of genes associated with both pro- and anti-inflammatory programs [97]. Most highly upregulated transcriptional regulator in AdoMacs was *HEY1* (**Fig. 2c**), a canonical Notch target and transcriptional repressor reported to attenuate secretion of inflammatory cytokines in macrophages [98]. Next, we inferred candidate upstream transcription factors (TFs) from observed transcriptional changes using TRRUST annotations (Suppl. Fig. 7a, b, Suppl. Table 5) [99]. Complementary TF binding site (TFBS) analysis was performed using the TFBS enrichment tool from the UniBind repository (Suppl. Fig. 7c, d, Suppl. Table 6) [100]. Consistent with the above observations, transcriptional changes in AdoMacs indicated increased activity of both the inflammatory TFs *NFKB1* and *RELA*, as well as anti-inflammatory TFs *STAT3* and *TFAP2A* (Suppl. Fig. 3a) [87, 101, 102]. TFBS analysis further indicated that promotor regions of DEGs in AdoMacs were enriched for binding sites of *STAT6*, a prominent driver of anti-inflammatory macrophage polarization (Suppl. Fig. 3c) [103].

Consistent with immunosuppressive pathway enrichment, we observed the upregulation of receptor genes linked to impaired macrophage-mediated cancer cell phagocytosis (*SIGLEC10, LILRB1*) and immunosuppressive TAM activation *(LAIR1, CD163, CD300E*) [5, 6, 88, 104, 105]. Importantly, we observed the downregulation of genes involved in MHC-II antigen presentation (*HLA-DMA/B, -DOA, -DPA1*) (**Fig. 2b**), together with their master transcriptional transactivator *CIITA* (log2FC < - 0.99, FDR < 0.015, Suppl. Table 4) [106]. Inference of TF activities also highlighted significantly reduced activity of the crucial transcriptional regulators of the antigen presentation pathways in the RFX family (*RFXANK, RFXAP, RFX5*, Suppl. Fig. 3b) [106]. This finding is particularly relevant in light of previous studies linking MHC-II^low^ TAM states to hypoxic tumor regions [107]. Furthermore, we observed the upregulation of the gene set “Tryptophan catabolism” (p_adj_ < 0.015) (Suppl. Table 2) including genes *IDO1* (**Fig. 2b**), *KYNU* and *KMO* from the kynurenine pathway, which represents the primary metabolic route for Trp degradation. Depletion of extracellular Trp can lead to the metabolic deprivation of T cells [7, 8]. Of note, hypoxia induces *IDO1* upregulation in dendritic cells and macrophages cocultured with hypoxic cancer cells upregulate *IDO1* gene [108, 109]. Among the few most highly upregulated genes in AdoMacs were TAM markers *MARCO* and *CHI3L2* (**Fig. 2b**) [110].

In addition to transcriptomic profiling, we performed proteomic and phosphoproteomic analyses of the macrophage states which displayed pronounced transcriptomic responses. Measurements were performed after 90 minutes of stimulation to assess early signaling cascade events. As expected at this early time point, total protein abundance showed only modest condition-specific changes, with most variation attributable to donor effects (Suppl. Fig. 9). Therefore, we normalized phosphorylation levels to total protein abundance (see Methods) and performed differential phosphorylation analysis (Suppl. Table 8) followed by the kinase enrichment analysis (KEA) (Suppl. Table 9). Among the proteins with differentially phosphorylated residues in AdoMacs compared to untreated macrophages (log2FC ≥ 1, FDR ≤ 0.05) (Suppl. Fig. 10a), several have previously been implicated in immunosuppressive TAM roles, such as LRRFIP1, IQGAP3 and SPP1 [111-113]. Additionally, we observed increased phosphorylation of the purinergic receptor P2RY11, a receptor for extracellular ATP, which has been linked to increased *IDO1* expression in dendritic cells and anti-inflammatory activity in macrophages [114, 115]. KEA further indicated increased activity of PDHK1 kinase, previously reported to be induced in macrophages during hypoxia [116], as well as increased activity of NEK7 kinase, implicated in the regulation of inflammation (**Fig. 2d**) [117].

Overall, these results indicate that Ado initiates a complex transcriptional program, which regulates inflammation, promotes Trp catabolism, and limits antigen presentation and tumor cell phagocytosis. Treatment with K^+^, which similarly to Ado accumulates in necrotic TME regions, initiated upregulation of several genes related to antiviral response and IFNα/β signaling (**Fig. 3a**, Suppl. Fig. 3e). This was illustrated by the upregulation of *IFIT1, IFIT2, IFIT3, IFIT5, IFIH1, ISG15,OAS1, OAS2, OAS3, DDX60* and *DDX58* genes (**Fig. 3b**) from said pathways, as well as the prominent upregulation of the regulatory TFs *IRF7* (log2FC > 2, FDR <0.001) and *STAT1* (log2FC > 1.4, FDR <0.005) (**Fig. 3c**) [118, 119]. Among most highly upregulated individual genes were *CCL24* and *CCR2* (**Fig. 3b**), which were both previously reported as markers for immuno-suppressive TAMs in a state with M2-like polarization [86, 120-122].

Similarly to AdoMacs, K^+^-stimulated macrophages (K^+^Macs) strongly downregulated antigen presentation genes *HLA-DQA1*, -*DOA, -DQB1*, -*DPA1* and their upstream transcriptional regulator *CIITA* (log2FC < -0.99, FDR <0.03) (**Fig. 3b, c**), as well as decreased inferred activity of TFs in the RFX family (*RFXANK, RFXAP, RFX5*) (Suppl. Fig. 7b) [106]. Importantly, we did not observe analogous shift towards downregulation of antigen presentation programs in M2a- or M2cMacs, thus highlighting the relevance of Ado- and K^+^Macs for recapitulating distinct functional aspects of clinical macrophage states.

Phosphoproteomics analysis revealed the upregulated phosphorylation of DDX41 (Suppl. Fig. 10b), a protein capable of initiating type I IFN response [123]. Additionally, we observed the upregulated phosphorylation of the SRC kinase, consistent with its reported roles macrophage function and antiviral immunity [124, 125] and its ability to block K^+^ efflux by inhibiting voltage-gated K^+^ channels [126, 127]. KEA analysis of upregulated phosphosites in K^+^Macs suggested increased activity of kinases SNRK, MST1, MEKK2 and TAK1 (**Fig. 3b**). SNRK is a key immunometabolic regulator, with reported roles in suppressing inflammation and promoting angiogenesis [128, 129], while MST1 has been previously implicated in macrophage polarization and oxidative stress responses [130, 131]. MEKK2 and TAK1 can phosphorylate and activate the IRF3 TF [132-134], a central regulator of the IFN I response that has been implicated in the inhibition of NFκB-dependent inflammatory signaling pathways [123, 135].

From a TME niche perspective, AdoMacs and K^+^Macs may both represent macrophage states shaped by metabolically stressed, hypoxic, or necrotic tumor regions, where extracellular Ado and potassium can accumulate. In line with this, our multi-omics analysis showed that both stimuli impaired antigen presentation, a feature previously associated with TAMs residing in hypoxic tumor regions [107, 136]. However, they differed in their dominant molecular programs: AdoMacs displayed a metabolically rewired phenotype, marked by MT expression, Trp catabolism, and concurrent inflammatory and immunosuppressive signaling, whereas K^+^Macs showed a more focused type I IFN-associated response.

### Distinct stimuli drive macrophage lipid metabolic rewiring

Cellular and molecular changes induced with TGFβ have been extensively studied. However, the specific effects on primary human macrophages on different molecular levels have so far not been characterized in depth. As expected, we observed here that TGFβ-stimulated macrophages (TGFβMacs) strongly upregulated TGFβ-, BMP- and SMAD-signaling pathways, as illustrated by the upregulation of *SMAD6* and *SMAD7* (**Fig. 4a**), consistent with known TGFβ signaling mechanisms [54]. Additionally, we observed significant upregulation of pathways associated with the ECM organization (**Fig. 4a**) as illustrated by the upregulated expression of matrix proteases (e.g. *ADAM12*), integrins (e.g. *ITGAV*) and other ECM components (e.g. *COL7A1*) (**Fig. 4b**). Downregulated gene sets indicated reduced interactions with lymphoid cells (Suppl. Table 2), as illustrated with downregulation of the immunostimulatory receptor CD169 (encoded by *SIGLEC1*) [137-139]. GSEA of hallmark genes additionally revealed a trend toward downregulation of IFNα/γ responses (Suppl. Fig. 3f). Among upregulated genes were the lipoprotein receptor gene *OLR1* (**Fig. 4b**) [140, 141], *PMEPA1* – a regulator of TGFβ-signaling [142], *AXL* – a receptor tyrosine kinase linked to immunosuppressive macrophage polarization [143] and abovementioned *APOC1*.

**Fig. 4:**
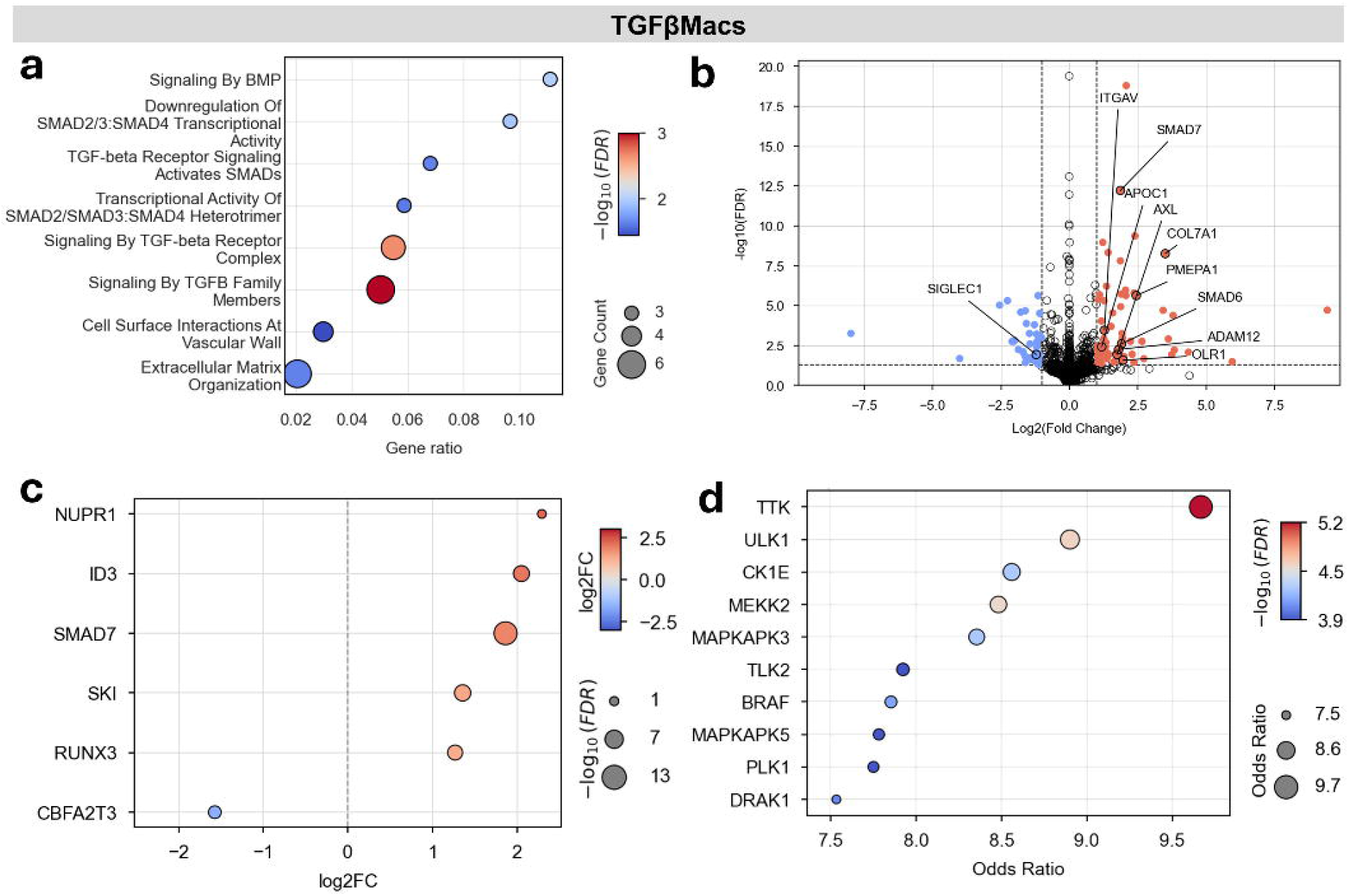
Characterization of the macrophage molecular responses to TGFβ. **a**, Dot plot depicting top over-represented Reactome pathways among DEGs in macrophages stimulated with TGFβ (TGFβMacs). DEGs are defined in comparison to untreated macrophages with the same thresholds as detailed in Figure 1. **b**, Volcano plot illustrating gene expression changes in TGFβMacs when compared to untreated macrophages. Graph axes, colors and thresholds are as in Figure 1. **c**, Dot plot illustrating top differentially up- and downregulated transcription factor genes in TGFβMacs when the gene expression levels are compared to untreated macrophages. A threshold of log2FC ≥ |0.5| and an FDR ≤ 0.05 were used to indicate significant TFs. **d**, Dot plot depicting kinases that are inferred to be upstream regulators of the significantly differentially upregulated phosphopeptides in TGFβMacs.

The most highly upregulated TF in TGFβMacs was *NUPR1* (log2FC > 2.2, FDR < 0.02) (**Fig. 4c**), which also has a role in immunosuppressive macrophage polarization [144]. Additionally, we observed a high differential expression of the TF *RUNX3* gene (log2FC > 1.2, FDR < 0.001), which has been previously linked to TGFβ-signaling and the suppression of inflammation [102]. Under the applied *in vitro* conditions, combined stimulation with TGFβ with IL-33 did not yield significant transcriptional changes when compared to the single stimulation with TGFβ (Suppl. Table 2).

Phosphoproteomics analysis after the TGFβ treatment revealed an increased phosphorylation of the SRC kinase, which has been reported to be activated by TGFβ receptors and necessary for downstream TGFβ-signaling [145] as well as the upregulated phosphorylation of ENO1 (Suppl. Fig. 10c, log2FC > 5.6, FDR < 0.001) a glycolytic enzyme that requires TGFβ/Smad3 signaling for membrane translocation and that has been previously proposed as a target in macrophage reprogramming [146]. We also observed the differential phosphorylation of the TF EGR2, which has been recently implicated in protumoral TAM functions [147]. KEA analysis revealed the potential involvement of upstream kinases previously described as regulators of TGFβ-signaling such as MEKK2, TAK1 and BMPR2 (**Fig. 4d**, Suppl. Table 9) [148]. Other single stimuli applied under the described conditions did not initiate pronounced coordinated changes in the expression levels of functionally related genes. However, the most differentially regulated genes nonetheless highlighted induction of relevant cellular processes. For instance, macrophages stimulated with Lac (LacMacs) – an immunosuppressive metabolite accumulating in the TME due to the Warburg effect [149]-upregulated pathways related to lipid metabolism, exemplified by the differential expression of *APOC1* and *ABCG1* genes (Suppl. Fig 6a, Suppl. Table 1). These genes have been previously reported to be linked to metastasis and T cell suppression when expressed by TAMs [34, 150, 151]. In addition, lactate stimulation induced a strong upregulation of TGFβ signaling in macrophages in Hallmark GSEA analysis (Normalized Enrichment Score (NES) > 1.8, FDR < 0.005, Suppl. Fig. 4a, Suppl. Table 3). Macrophages stimulated with oleate differentially upregulated the *PDK4* gene - a metabolic switch favoring fatty acid oxidation, and the *PLIN2* gene - a marker of lipid-associated macrophages involved in lipid storage and EMT, (Suppl. Table 1) [152, 153]. Oleate is an unsaturated fatty acid abundant in the TME, which was in previous studies shown to be able to promote M2-like macrophage polarization and lipid droplet formation [67]. Jointly, these results suggest that independent stimuli reshape macrophage metabolism while simultaneously affecting additional programs, such as in the case of TGFβ, regulating lipid metabolism while promoting ECM remodeling and suppressing inflammatory pathways.

### Individual stimulus signatures persist in tumor-conditioned medium-treated macrophages

Next, we sought to compare molecular changes in macrophages activated with single stimuli to those exposed to a complex mixture of cytokines and metabolites from TCMs. We obtained TCMs from the fresh primary brain tumor resections: three of them being classified as meningiomas and one as vestibular schwannoma (see Methods). First, we analyzed the composition of TCMs using a cytokine array and ELISAs, providing insight into present cytokines, but not metabolites. This revealed patient-specific composition, but also confirmed the shared presence of signaling mediators that have been previously reported to affect macrophage function, such as VEGF-A, IL-33, and LIF (Suppl. Fig. 11) [69, 70, 75, 78]. Next, we exposed differentiated macrophages to TCMs following the same protocol as above. This analysis revealed heterogeneous transcriptional phenotypes across treated macrophages, with the TCM containing the highest level of the total detected cytokines (TCM2) also inducing the strongest transcriptional response (**Fig. 1c**). Across TCMs, macrophages converged on a shared immunosuppressive baseline, while individual patient samples differed in the strength and combination of conserved programs, generating two dominant response archetypes: an IFN-like state and a metabolic/TGFβ-like state.

We found that macrophages exposed to either of the four different patient TCMs upregulated pathways associated with general cytokine- and IL-10 signaling pathways and displayed mixed functional phenotypes (Suppl. Fig. 5). Furthermore, three out of four analyzed TCMs (TCM2, TCM3 and TCM4) initiated upregulation of the genes associated with the GO term ‘axon interactions’, thus reflecting the presence of the host tissue components in the TCMs (Suppl. Table 2). Furthermore, two conditions (TCM2 and TCM4) upregulated genes from the pro-angiogenic pathways (Suppl. Table 2). Of note, in macrophages stimulated with TCM1 we observed a similar transcriptional program as in K^+^Macs: the upregulation of IFNα/β signaling components (Suppl. Fig. 4b, 5a), such as *DDX58, ISG15, IFIT1, IFIT2, IFIT3, IFIT5, OASL, OAS1, OAS2, OAS3* and *IRF7* genes, with a parallel significant downregulation of MHC-II antigen presentation gene *HLA-DQA1* (Suppl. Table 1, 2). Interestingly, macrophages stimulated with TCM2, TCM3 and TCM4 showed prominent upregulation of the TGFβ signaling pathway genes (Suppl. Fig. 4c-e), despite only low concentrations of TGFβ being detected in the respective media (Suppl. Fig. 11). In addition, LacMacs and macrophages stimulated with TCM2, TCM3 and TCM4 showed a substantial overlap in differentially upregulated genes. Lac induced statistically significant upregulation of 34 genes, 28 of which were also upregulated in macrophages exposed to TCM2, TCM3, and TCM4 (Suppl. Table 10). Considering that Lac stimulation alone was sufficient to trigger a significant TGFβ signaling response in the Hallmark GSEA analysis (NES > 1.8, FDR < 0.01, Suppl. Fig. 4a, Supplementary Table 3), it is plausible that TGFβ-transcriptional signatures can potentially arise from multiple factors in the TME, including Lac. Additionally, all TCM-stimulated macrophages differentially expressed the *PDK4* gene, and macrophages stimulated with TCM2, TCM3 and TCM4 expressed *PLIN2* gene. In this study, both genes were exclusively induced by oleate.

**Fig. 5:**
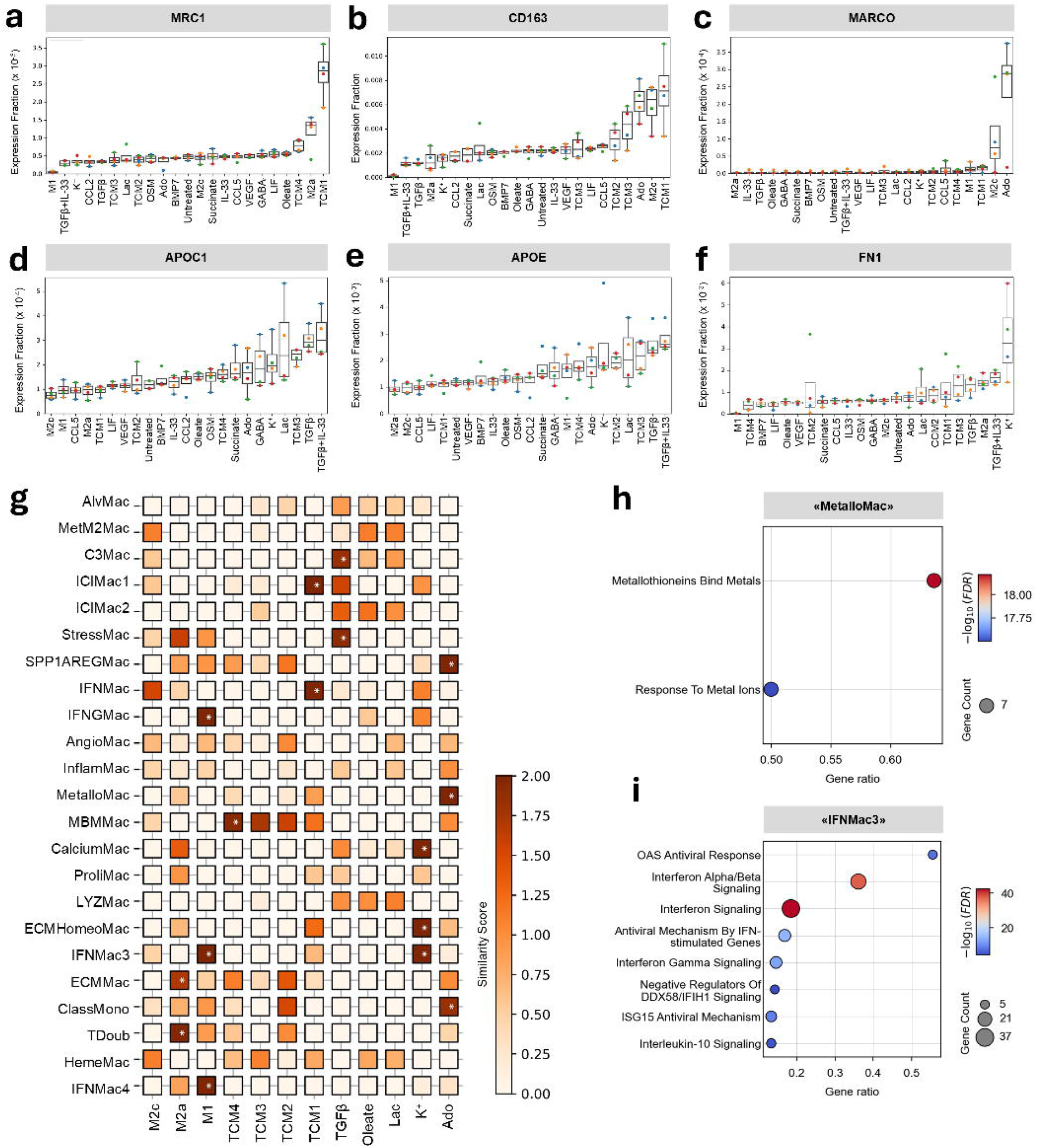
*In vitro* macrophages share similarities with clinically relevant *in vivo* macrophage states. **a-f**, Boxplots illustrating expression levels of TAM markers in *in vitro* generated macrophage states. Boxplots indicating median, inter quartile ranges as well as individual values across donors are shown. **a**, MRC1. **b**, CD163. **c**, MARCO. **d**, APOC1. **e**, APOE. **f**, FN1. **g**, Matrix depicting similarity scores between generated *in vitro* macrophage states and previously defined clusters of tumor-associated macrophages, reported in the pan-cancer patient scRNA-seq study by Coulton et al. [21], are shown. Similarity scores indicate the magnitude of the second-order distance shift of the treated state relative to the untreated control; P values are obtained through the statistical comparison between untreated and treated conditions via permutation testing following with Bonferroni correction; *indicates a *p* value < 0.05. **h-i**, Dot plots depicting over-represented Reactome pathways among differentially upregulated genes (avg_log2FC ≥ 0.5) in the clusters of patient tumor-associated macrophages which were defined as **h**, “MetalloMac” and i, “IFNMac3” in the study by Coulton et al. [21].

**Fig. 6:**
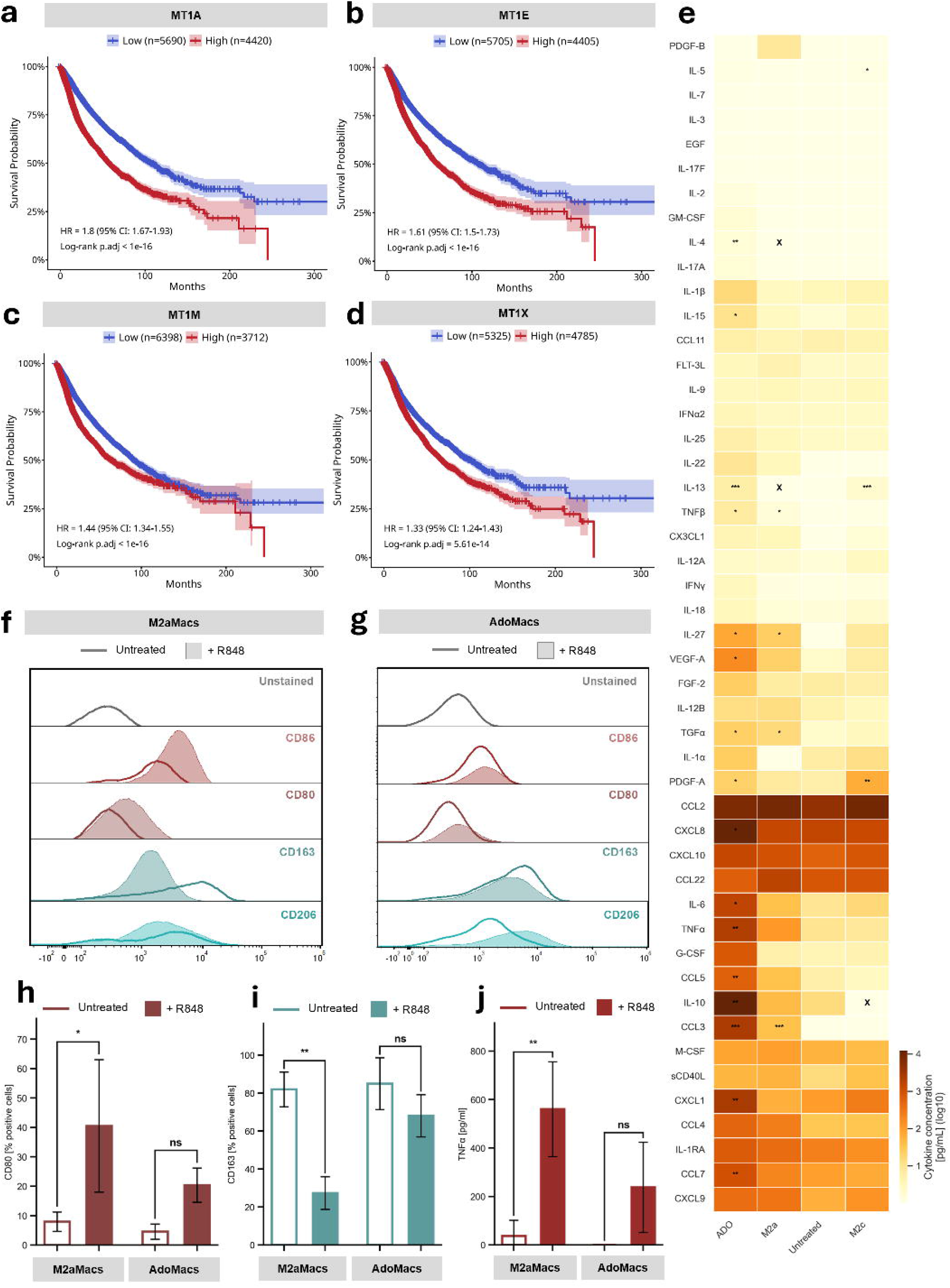
Adenosine-stimulated macrophages model clinically relevant mechanisms. **a-d**, Kaplan-Meier curves illustrating the association of metallothionein gene expression levels with the patient survival in pan-cancer TCGA-cohorts. **a**, MT1A. **b**, MT1E. **c**, MT1M. **d**, MT1X. **e**, Heatmap illustrating cytokine concentrations measured in the supernatants of M2a-, M2c-, AdoMacs and untreated macrophages. Color gradient represents log_10_(cytokine concentration [pg/mL]). For all conditions, average values across 3 biological replicates are shown; Fields marked with X indicate exclusion due to analyte-stimulus overlap. Statistical comparison between untreated vs. treated conditions was performed with one-way ANOVA followed with FDR correction; \**p* < 0.05, \*\**p* < 0.01, \*\*\**p* < 0.001. **f-g**, Representative flow cytometry histogram plots of polarized macrophages states with and without R848 treatment. Values for the established macrophage surface markers CD86, CD80, CD163 and CD206 are shown. Figure **f**, shows M2aMacs. And figure g, the same for AdoMacs. **h-i**, Bar charts illustrating the percentage of cells positive for surface markers assessed through flow cytometry for M2a- and AdoMacs with and without R848 treatment. Figure h, shows CD163 levels, Figure **i**, CD80 levels. Both graphs represent values from 3 biological replicates; error bars indicate standard deviation, P values are calculated with a one-way ANOVA with Tukey’s multiple comparisons test; \**p* < 0.05, \*\**p* < 0.01, \*\*\**p* < 0.001. **j**, Bar chart illustrating the secreted TNFα concentration of M2a- and AdoMacs with and without R848 treatment. n = 3 biological replicates; error bars indicate standard deviation; P values are calculated with a oneway ANOVA with Tukey’s multiple comparisons test; \**p* < 0.05, \*\**p* < 0.01, \*\*\**p* < 0.001.

Collectively, transcriptomic analysis of distinct macrophage molecular states showed that signaling routes induced by individual TME-relevant stimuli were recapitulated in macrophages exposed to patient-derived TCMs.

### *In vitro* generated macrophages mimic features of clinically relevant TAM subsets

To assess the clinical relevance of macrophage states generated *in vitro*, we first examined a panel of TAM markers previously associated with immunosuppressive, metabolic, or pro-tumoral programs in patients. The canonical immunosuppressive M2-like markers CD206 (encoded by *MRC1*) and CD163 were differentially induced across conditions: *MRC1* was upregulated in M2aMacs, as well as macrophages stimulated with TCM1 and TCM4, whereas *CD163* was highly expressed in M2cMacs, TCM-stimulated macrophages (TCM1, TCM2, TCM4) and AdoMacs (**Fig. 5a, b**). Among TAM-associated markers, *MARCO* was induced in M2cMacs, but more prominently in AdoMacs (**Fig. 5c**). The lipid-associated TAM markers *APOE* and *APOC1* were highly expressed in macrophages stimulated with TGFβ, Lac, and TCM3 (**Fig. 5d, e**). Notably, *APOE* expression in TAMs has been previously linked to TGFβ signaling [33] and, as discussed above, TGFβMacs, LacMacs, and TCM3Macs all exhibited upregulation of TGFβ signaling components (Suppl. Fig. 3f, 4a, d). Finally, *FN1* expression in TAMs has previously been linked to metastasis, recurrence and immunosuppression [36, 37], and was here upregulated in K^+^Macs by high extracellular K^+^ exposure (**Fig. 5f**).

To further contextualize the physiological relevance of *in vitro* generated macrophage states beyond individual markers expression, we aligned their transcriptomic profiles with TAM clusters reported in a recently published pan-cancer TAM atlas comprising > 300.000 scRNA-seq TAM signatures from > 30 patient studies [21]. This atlas identified recurrent TAM phenotypes across tumor types, including inflammatory (e.g. “IFNGMac”, “IFNMac3”, “IFNMac4”), opsonization-associated (“C3Mac”), MT-expressing (“MetalloMac”), ECM-remodeling (“ECMMac”, “ECMHomeoMac”), and melanoma brain metastasis-enriched TAM clusters (“MBMMac”). We compared our bulk RNAseq profiles with these defined TAM clusters by calculating treatment-induced expression changes as the magnitude of second-order distance shifts relative to untreated macrophages. The resulting scores were z-normalized per condition and cluster, and statistical significance was assessed by permutation testing (see Methods). When aligned with the patient TAM atlas (**Fig. 5g**), M1Macs showed significant similarity with inflammatory TAM clusters “IFNGMac”, “IFNMac3” and “IFNMac4” (FDR <0.03). The similarity was primarily driven by the high expression of inflammatory genes such as *CXCL10, MX1*, and *IFIT3*. M2aMacs significantly mapped to cluster “ECMMac” (FDR < 0.05), a TAM state linked to ECM remodeling [21]. The observations are consistent with the known roles of M1- and M2-like states. Furthermore, AdoMacs displayed high similarity with the cluster “SPP1AREGMac” (FDR < 0.02), driven by the upregulation of *SOD2*, which has a role in the protection against oxidative stress [154, 155], as well as *IL1B, VCAN, CXCL8* and *S100A/9*, which have roles in angiogenesis [156-159]. In addition, the AdoMacs showed a striking similarity with the “MetalloMacs” cluster (FDR < 0.02), driven by the strongly upregulated expression of MT genes (*MT1A, MT1X, MT2A, MT1E, MT1H, MT1F*, and *MT1M*). ORA of upregulated genes in Coulton et al. [21] clusters identified enrichment of MT-associated pathways in the “MetalloMac” cluster. (**Fig. 5h**), mirroring the enrichment observed in AdoMacs. Of note, MT genes expressed exclusively in *in vitro* AdoMacs were also highly expressed exclusively in the clinical TAM cluster MetalloMacs (Suppl. Fig. 12). Moreover, MetalloMacs displayed downregulation of pathways associated with MHC-II antigen presentation, equivalently to AdoMacs. K^+^Macs aligned significantly with the “IFNMac3” cluster (FDR < 0.02), sharing high expression of the canonical genes of a type I IFN response such as *MXs, IFITs* and *STAT1*, and downregulation of MHC-II antigen presentation genes *HLA-DQB1* and *HLA-DQA1*. ORA of genes upregulated in the “IFNMac3” cluster showed substantial overlap with pathways induced in K^+^Macs, with both clusters showing overrepresentation of pathways related to antiviral response and IFNα/β signaling (**Fig. 5i**). K^+^Macs also showed a significant similarity with the TAM clusters “ECMHomeoMac” (FDR < 0.03) and “CalciumMac” (FDR < 0.02) with which they shared the high expression of *FN1*. TGFβMacs aligned significantly with the TAM cluster “C3Mac” (FDR < 0.04), driven by the upregulation of the *C3* complement gene, as well as high expression of *APOC1* and *NUPR1*, both associated with immunotherapy resistance when expressed by TAMs [35, 144]. Finally, all states stimulated with TCMs showed a strong similarity to the cluster “MBMMac” associated with melanoma brain metastasis, likely reflecting the effect of the host organ factors as the TCMs used here were derived from primary patient brain cancer tissues. Overall, these results demonstrate that multiple *in vitro* generated macrophage states recapitulate prominent features of TAM phenotypes identified in patients, suggesting their relevance for understanding diverse aspects of physiological TAM functions.

### AdoMacs exhibit TAM-like features linked to poor prognosis

Motivated by the observation that *in vitro* AdoMacs recapitulate prominent features of the clinical TAM cluster “MetalloMac”, we evaluated the prognostic impact of the highly upregulated MT genes. For this, we analyzed survival data from the TCGA dataset and performed Kaplan-Meier survival analysis. We individually analyzed 11 MT genes prioritized from the differential expression analysis in AdoMacs. In a pan-cancer analysis, higher expression levels of *MT1A, MT1E, MT1M, MT1X, MT1DP, MT1L, MT2A and MT1JP* were significantly associated with decreased overall survival (OS) after multiple testing correction with hazard ratios (HRs) ranging from 1.2 to 1.8 (**Fig. 6a-d**, Suppl. Fig. 14).

Ado has been reported to induce immunosuppressive macrophage polarization independent of interleukin-4 receptor signaling, resulting in the increased secretion of IL-10, VEGF-A and other anti-inflammatory mediators [49-51]. On the transcriptome level, we observed upregulation of the IL-10 pathway in AdoMacs (**Fig. 2a**), but not a significant overexpression of the *IL10* and *VEGFA* genes (Suppl. Table 1). To investigate this further, we performed a cytokine array on cell culture supernatants from AdoMacs obtained after 24 h of polarization and compared them to supernatants of untreated macrophages and canonical M2 *in vitro* phenotypes (M2a- and M2cMacs) (see Methods). AdoMacs indeed secreted significantly increased levels of IL-10 (FDR < 0.003) and VEGF-A (FDR < 0.05) compared to untreated macrophages (**Fig. 6e**), exceeding concentrations observed in M2a- and M2cMacs. In addition, AdoMacs secreted elevated levels of IL-4 (FDR < 0.004) and IL-13 (FDR < 0.001), which could further reinforce immunosuppressive loops in the TME. Additionally, compared to the other states, AdoMacs released significantly elevated levels of CCL7, CXCL1, CCL3, CCL5, TNFα/β, CXCL8, IL6, PDGF-A, TGFα, IL-15 and IL-27 indicating a broader activation spectrum than previously reported. Collectively, these results suggest that elevated Ado levels in the TME, typically located in the tumor necrotic core, could contribute to the polarization of macrophage states that support tumor progression and link to poor clinical outcomes.

Macrophages respond to new stimuli depending on their initial polarization state [160, 161], adding complexity to studying their behavior in the dynamic TME and complicating therapeutic macrophage□reprogramming efforts. TLR-agonist R848 is known to reprogram canonical M2a-like macrophages stimulated with IL-4 and IL-13 toward a more inflammatory phenotype *in vitro* [44, 162], however this has so far not been demonstrated with other macrophage populations. Thus, we evaluated the reprogramming efficiency of R848 on here generated macrophage states. For this, we treated the pre-polarized macrophage states with R848 for 24 h and assessed the resulting changes in the TNFα secretion with ELISA and changes in the levels of representative surface receptors with a flow cytometry. As expected, we observed that R848 resulted in a pronounced reprogramming response in M2aMacs, manifested in a significant reduction of the M2-like surface receptor CD163 levels and a significant increase of the M1-like CD80 receptor levels as well as high increase in the TNFα secretion (**Fig. 6f, h-j**). However, in AdoMacs, R848 treatment could not induce an equally significant inflammatory phenotypic shift, indicating that AdoMacs may be less susceptible to the drug than M2aMacs (Fig. 6g-j). Overall, we found that here generated macrophage states differed substantially in their magnitude in phenotypic shifts in response to R848 (Suppl. Fig. 15), underscoring the need for macrophage-state-specific reprogramming strategies rather than a one-size-fits-all approach.

## DISCUSSION

Macrophages in the TME are exposed to a highly complex and dynamic stimuli and the exact recapitulation of the TAM states *in vitro* is likely not possible. However, recent systematic analyses of immune cell responses to individual cytokines [59, 60] have provided unprecedented insights into the plasticity of the transcriptional responses of these cells. Here, we found that *in vitro* macrophage states induced through a stimulation of primary human macrophages with defined TME-derived factors shared molecular similarities with clinically relevant TAM populations. Transcriptomic and phosphoproteomic datasets generated here for individual states hence provide mechanistic insights into how TME-derived stimuli may contribute to the emergence of tolerogenic and immunosuppressive macrophage functions in tumors.

For several stimulus-induced states, these findings expand our knowledge from prior studies conducted in mouse models or studies focused on the assessment of a limited number of cellular markers. For instance, elevated extracellular K^+^ is reported to induce strongly immunosuppressive polarization of mouse macrophages and act as a metabolic and transcriptional switch when supplemented to TCM, which could be mitigated by inhibiting K^+^ influx [48]. Together, these findings suggest that extracellular K^+^ does not directly induce a canonical immunosuppressive macrophage state, but may instead rewire inflammatory responsiveness by downregulating MHC-II antigen-presentation genes while skewing macrophages toward a type I IFN-associated program, which can repress inflammasome activation [163], potentially acting synergistically with other immunosuppressive factors in the TME.

Earlier studies characterized Ado as an M2-like driver of macrophage polarization, based on increased IL-10 and VEGF expression and secretion identified using PCR and cytokine quantification assays [49-51]. Our data are consistent with these observations, but extend them by showing that Ado regulates a far broader spectrum of potentially tolerogenic pathways, including the upregulation of Trp catabolism which could contribute to the metabolic deprivation of T cells, the upregulation of myeloid checkpoint genes which can inhibit tumor cell phagocytosis, the transition towards an MHC^low^ phenotype and the distinct upregulation of MTs. Although MT-expressing TAMs have been reported in multiple scRNA-seq studies of patient tumors [21, 23, 90], their role within the TME is yet to be fully understood. By recapitulating the MT-expressing phenotype *in vitro* through Ado stimulation, we establish an experimental framework to further investigate the functional aspects of this state. This is of particular interest, as we observed here that the upregulation of several MT genes in the patient tumor samples associated with poor survival. However, because this survival analysis is based on bulk tumor transcriptomes, the cellular source of MT expression cannot be assigned specifically to macrophages. Nevertheless, the functional relevance of AdoMacs is supported by its reduced responsiveness to R848-mediated reprogramming compared with canonical M2aMacs, suggesting that Ado-conditioned macrophages may represent a particularly difficult-to-reprogram TAM-like phenotype. Prior work showed that Ado enhances IL-10 secretion primarily through Ado-receptor signaling and post-transcriptional regulation rather than transcriptional induction [50]. We also observe the mismatch between quantified cytokine secretion and IL-10 mRNA levels in our dataset, highlighting inherent limitations of scRNA-seq to fully resolve all relevant aspects of functional TAM states. Of note, Ado has been previously reported to upregulate *IL6, CXCL8, IDO1* and *ARG2* in dendritic cells, and our results confirm that this also occurs in macrophages [108, 164]. Ado and K^+^ are abundant in tumor necrotic cores. Future spatial transcriptomics studies may help determine whether TAMs with high expression levels of MTs and *FN1* genes are preferentially localized to these regions.

Next, TGFβMacs represent a highly studied macrophage phenotype. However, there is only a limited number of studies on primary human macrophages [165]. On the transcriptome level, TGFβMacs demonstrated a strong increase in the expression levels of genes with roles in ECM remodeling, and an increased expression of lipid metabolism–associated TAM markers *APOE* and *APOC1*, which are of increasing interest as therapeutic targets [33, 35]. Notably, LacMacs and certain TCM-stimulated macrophages also exhibited TGFβ-signaling signatures, indicating convergence of multiple TME-derived cues on TGFβ-like transcriptional programs.

The observed similarities in comparison to a pan-cancer TAM atlas [21] support the notion that discrete TME-derived factors can drive key programs that play a role in clinically relevant macrophage states. However, even a high transcriptional similarity to a given TAM cluster reflects only partial recapitulation of co-occurring gene programs rather than full reproduction of an entire *in vivo* state. Furthermore, several stimuli in our study, which had demonstrated roles in macrophage polarization in past studies, did not elicit significant transcriptional changes here, which may also indicate the need for higher concentrations or longer stimulation periods. The same limitations apply for time-dependent phosphorylation dynamics in response to applied stimuli. Finally, donor variability remains an inherent challenge in primary immune cell studies [60]; although strong stimuli produced reproducible shifts across donors, larger cohorts will be necessary to fully map genetic backgrounds on macrophage plasticity.

Overall, this work provides systematic molecular context for clinically relevant macrophage markers and illustrates how TME-derived cues can shape human macrophage polarization and reveal stimulus-linked programs that contribute to clinically relevant TAM heterogeneity. It further provides an experimental framework for replicating aspects of clinically relevant signaling programs *in vitro* and thus offering a critical tool for the design of TAM-state-specific therapeutic strategies.

## METHODS

### Experimental section

#### Generation of tumor-conditioned medium

The protocol for the generation of TCM was adapted from Jacob, Salinas et al [166]. After ethical approval (BASEC Nr. 2021-00473), fresh surgically resected tissue from central nervous system neoplasms of four patients (Table 1) was placed in Hibernate A medium (BrainBits) and kept at 4 °C or transported on ice. The tissue was transferred to a sterile glass dish, washed with sterile DPBS and submerged in RPMI-1640 medium containing 10 %FCS and 1 % PS. After removal of major blood vessels and cauterized areas, the tumor tissues were minced into pieces with a diameter of 0.5-1 mm using fine dissection scissors, a scalpel or a 1 mm biopsy puncher. The tumor pieces were incubated in sterile distilled water for 5 min to lyse contaminating red blood cells through osmotic shock, before being distributed in 6-well plates containing 4 mL of RPMI-1640 medium containing 10 %FCS and 1 % PS. The plates were placed on an orbital shaker rotating at 120 rpm within a 37 °C, 5 % CO_2_, and 90% humidity sterile incubator for 3 days. The generated TCM supernatant was removed and sterile filtered. TCMs were stored at -80 °C until further use. The cytokine composition of TCMs was analyzed using the Human Cytokine 96-Plex Discovery Assay (EveTechnologies), as well as ELISA kits (ThermoFisher) for IFNγ, IL-4 and TGFβ.

### In vitro generation of macrophage subsets

Buffy coats from four healthy human donors were purchased from the Zurich Blood Bank (#D9993V00) after ethical clearance (BASEC Nr. 2021-00687) and project approval and purified using a density gradient centrifugation with Ficoll (Sigma-Aldrich). The monocytes were isolated from the peripheral blood mononuclear cells (PBMCs) using a CD14 magnetic bead (Miltenyi Biotech) selection. After differentiation with 15 ng/mL GMCSF for M1Macs and 30 ng/mL M-CSF for all other subsets for 6 days at 37 °C in RPMI-1640 medium (Sigma-Aldrich) containing 10 % fetal calf serum (FCS) and 1 % penicillin/streptomycin (PS), the medium was replaced with polarization medium according to Table 2. Concentrations of 20 ng/mL were used for stimulation with cytokines, whereas metabolite concentrations were used that were previously reported to be relevant in the TME or for macrophage stimulation. For transcriptomics experiments, differentiated macrophages from 4 donors were polarized with their respective stimulus medium for 24 h before being washed with DPBS and incubated with cold harvesting solution (10 mM EDTA (Sigma-Aldrich) in DPBS) for 10 min at 4 °C. Cells were detached using a cell scraper (VWR). The cell pellet was resuspended in lysis buffer (Buffer RLT, Qiagen) containing 2 % Dithiothreitol (DTT) and homogenized using a QIAshredder (Qiagen) before isolating the RNA using the RNeasy Micro Kit (Qiagen) according to manufacturer’s instructions. The time point for the detection of early phosphorylation events was determined through Western Blot with a phospho-SMAD2/3 antibody (Invitrogen, RRID: AB_2855566) after stimulation with TGFβ for 0 min, 15 min, 30 min, 60 min, 90 min, 4 h, 6 h, 12 h, 24 h and 48 h, showing the highest phosphorylation rate at 90 min (Suppl. Fig. 4). For phosphoproteomics analysis, differentiated macrophages from 6 donors were therefore incubated with their respective stimulus solution for 90 min before being washed with DPBS and lysed in 200 μl Lysis buffer (2 % Sodium dodecyl sulfate (SDS) in 50 mM Tris/HCl pH 8.2) and snap-frozen in liquid nitrogen. All samples were transported on dry ice to the Functional Genomics Center Zurich.

#### Western blot

Differentiated macrophages were exposed to TGFβ for 0 min, 15 min, 30 min, 60 min, 90 min, 4 h, 6 h, 12 h, 24 h and 48 h. Cells were collected through scraping and centrifuges at 2000 rpm for 5 min, washed once with PBS, and lysed in RIPA buffer (50 mM Tris-HCl, pH 7.4, 150 mM NaCl, 1 mM EDTA, 1% NP-40, 0.25% sodium deoxycholate) supplemented freshly with protease inhibitor cocktail, 1 mM PMSF, and 1 mM DTT. Lysates were incubated on ice for 30 min and cleared by centrifugation at 13000 rpm for 10–20 min at 4°C. Supernatants were collected as whole-cell extracts, and protein concentration was determined by BCA assay. Equal amounts of protein (50 µg) were mixed with sample buffer containing DTT, heated at 95°C for 5 min, and resolved by SDS-PAGE on 4–12% Bis-Tris gels (NuPAGE, Invitrogen) in MOPS running buffer. Proteins were transferred to PVDF membranes using wet transfer at 100 V for 1 h in transfer buffer containing 20% methanol. Membranes were blocked in 5% BSA in PBS-T and incubated overnight at 4°C with primary antibodies (pSMAD2/3, Invitrogen, AB_2855566) diluted in PBS-T. After washing in PBS-T, membranes were incubated with HRP-conjugated secondary antibodies (Cytiva, RRID:AB_2650489) for 60 min at room temperature, washed again, and developed using SuperSignal West Dura substrate (Thermo Scientific). β-actin was used as a loading control (RRID: AB_2537667).

#### Macrophage reprogramming

Macrophage subsets were generated as discussed above from three biological replicates per condition. After 24 h of treatment with the polarization medium, cells were washed in DPBS and the medium was replaced with fresh culture medium containing 1 mg/mL Resiquimod (Sigma-Aldrich) and incubated for 24 h at 37 °C. Cells were washed with DPBS before being incubated at 4 °C for 10 minutes with harvesting solution. Subsequently, cells were detached using a cell scraper (VWR) and fixed using 4 % paraformaldehyde solution. Fixed cells underwent two blocking steps with 10 % FBS in DPBS for 1 h at 4 °C and 5 % Human Seroblock (BioRad) for 30 minutes at 4 °C. Subsequently, cells were stained using reaffinity antibodies for flow cytometry (Miltenyi Biotec) for human CD80 (RRID: AB_2751432), CD86 (RRID: AB_2727371), CD206 (AB_3663295) and CD163 (RRID: AB_2655475) for 30 minutes at 4 °C. Cells were washed and resuspended in FACS buffer (0.5% (w/v) Bovine Serum Albumin in DPBS). Cell culture supernatant of reprogrammed macrophage states were analyzed using a TNFα ELISA kit (ThermoFisher). Marker expression and cytokine secretion were analyzed by two-way ANOVA using Condition (e.g. M2a vs. ADO), Treatment (Control vs. R848). Post hoc comparisons were performed using Tukey’s multiple-comparisons test.

#### Cytokine quantification of in vitro generated macrophages

The cytokine composition of cell culture supernatants of generated macrophage states (stimulated with respective polarization medium for 24 h) was analyzed using the Human Cytokine 96-Plex Discovery Assay (EveTechnologies). Cytokine concentrations were measured in three independent biological replicates per condition. Conditions in which the analyte would be directly affected by the applied stimulus were excluded from the measurement (e.g. IL-10 quantification M2cMacs). For each cytokine, differences between treatment conditions (M2a-, M2c-, and AdoMacs) and the control (Untreated) were assessed by one-way ANOVA implemented as a linear model (*value* ∼ *condition*) after log2 transformation of the data. P-values for comparisons versus the untreated control were extracted from the same model and adjusted for multiple testing using the Benjamini–Hochberg false discovery rate method. FDR values below 0.05 were considered statistically significant.

### Transcriptomics analysis

#### Library preparation

The quality of the isolated RNA was determined with a Fragment Analyzer (Agilent, Santa Clara, California, USA). Only those samples with a 260 nm/280 nm ratio between 1.8–2.1 and a 28S/18S ratio within 1.5–2 were further processed. The Tecan® Universal Plus™ mRNA-Seq library preparation kit (Tecan Trading AG, Switzerland) was used in the succeeding steps. Briefly, total RNA samples (100-1000 ng) were poly A enriched and then reverse-transcribed into double-stranded cDNA. The cDNA samples were fragmented, end-repaired and adenylated before ligation of an anchor. Fragments containing the anchor on both ends were selectively enriched with PCR at the same time adding the index with UDI. The quality and quantity of the enriched libraries were validated using the Fragment Analyzer (Agilent, Santa Clara, California, USA). The product is a smear with an average fragment size of approximately 260 bp. The libraries were normalized to 10nM in Tris-Cl 10 mM, pH 8.5 with 0.1% Tween 20.

#### Cluster Generation and Sequencing

The NovaseqX (Illumina, Inc, California, USA) was used for cluster generation and sequencing according to standard protocol. Sequencing was paired end at 2 X150 bp.

#### RNA-seq bioinformatic analysis

The bioinformatic analysis pipeline was developed using Snakemake (version 9.9.0, RRID:SCR_003475) and can be found in the following repository: https://github.com/rriupu/mf_manuscript_transcriptomics_pipeline/tree/main.

#### Genomic data

The hg38 analysis set genomic sequence was obtained from the UCSC repository (https://hgdownload.soe.ucsc.edu/goldenPath/hg38/bigZips/analysisSet/hg38.analysisSet.fa.gz) [167]. Gene annotations in GTF format were also obtained from the UCSC repository (https://hgdownload.soe.ucsc.edu/goldenPath/hg38/bigZips/genes/hg38.ncbiRefSeq.gtf.gz).

#### Preprocessing and mapping

We removed adapter sequences from the raw reads using cutadapt (version 4.4) [168] together with the following parameters: --*minimum-length*=*20*, --*quality*-*cutoff*=20, --*nextseq*-*trim*=*20*, and --*poly*-*a*. We used STAR (version 2.7.0f, RRID:SCR_004463) [169] to index the genome and align the reads. Next, we used htseq-count (version 2.0.9) [170] to produce a gene count matrix with the following parameters: -*s yes*, -*r pos*, -*t exon*, and -*i gene*_*id*. In addition, we used FastQC (version 0.12.1, RRID:SCR_014583) and MultiQC (version 1.29, RRID:SCR_014982) [171] to run quality controls of raw, trimmed, and aligned reads.

#### Downstream analysis

All downstream statistical analyses were performed using R 4.4.1. We used the vst function from the DESeq2 package (version 1.46.0) [172] to perform a variance stabilization prior to the exploratory data analysis.

#### Exploratory data analysis

We explored the gene count data by plotting all pairwise sample-sample Euclidean distances and correlations. Together with a PCA, this highlighted that samples tended to cluster more by donor than by treatment, which lead to the inclusion of the donor information in the differential expression analysis. We used the removeBatchEffect function from the limma package (version 3.62.2, RRID:SCR_010943) [173] to remove the donor effect during the exploration of the data (Suppl. Fig. 1).

#### Differential expression analysis

We used DESeq2 (version 1.46.0, RRID:SCR_015687) [172] to run the differential expression analysis. After observing the donor effect during the exploratory data analysis, we included this variable in the analysis to measure the effects of the treatment while controlling for donor. We defined DEGs as log2(FoldChange) ≥ 1 and an FDR ≤ 0.05 and differentially expressed transcription factors as being expressed with log2(FoldChange) ≥ 0.5 and an FDR ≤ 0.05.

#### Transcription factor activity

TFs whose downstream targets were over-represented in the sets of differentially expressed genes were identified using TRRUST annotations [99]. We calculated a TF activity score

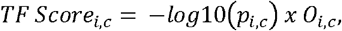

where *p*_*i,c*_ is the enrichment p-value for TF *i* in condition *c* and *O*_*i,c*_ is the number of overlapped genes predicted to be regulated by TF *i* in condition *c*, for each significantly activated TF (*p*_*i,c*_ ≤ 0.05 and *O*_*i,c*_ ≥ 3) in order to rank important TFs per condition.

#### UniBind Transcription Factor Binding Site Enrichment Analysis

We defined promoter regions of all protein-coding genes as -300 bp and + 100 bp from each gene’s transcription start site (TSS). TSS coordinates were obtained from the UCSC table browser [167]. We ran the TFBS enrichment analysis using the UniBind [100] TFBS enrichment analysis tool (https://bitbucket.org/CBGR/unibind_enrichment/src). We defined foreground regions as the TSSs of DEGs (FDR ≤ 0.05, log2FC> |2|). Additionally, we used the TSSs of all genes analyzed in the RNA-seq as the background. Finally, we used the precomputed LOLA database with the robust set of human TFBSs (https://zenodo.org/records/4704641).

#### Gene set enrichment and over-representation analysis

Differential expression results were subjected to preranked gene set enrichment analysis using GSEApy (RRID:SCR_025803) on each condition using MSigDB Hallmark gene sets [83]. Genes were ranked by signed FDR, calculated as −log_10_(FDR) × sign(log_2_FC). GSEA was run with 1000 permutations, a gene set size range of 15–500, and results were summarized using NES and FDR q-values across conditions. Over-representation analysis was performed using Reactome (RRID:SCR_003485) [81, 174] and Gene Ontology Biological Process (GO BP, RRID:SCR_002811) [82, 175] gene set libraries accessed via the Enrichr API through the GSEApy Python package. Analyses were restricted to human gene sets (Reactome_2022; GO_Biological_Process_2021). Only DEGs with a log2(FoldChange) ≥ 1 and an FDR ≤ 0.05 were considered in the over-representation analysis. Pathways and functional terms identified as being enriched with an adjusted p-value threshold of ≤ 0.05 were considered significant. Equivalently, ORA of differentially upregulated genes in TAM clusters identified in Coulton data [21] was performed using Reactome using a threshold of avg_log2FC≥ 0.5, and avg_log2FC≤-0.5 for differentially downregulated genes.

### Proteomics and phosphoproteomics analysis

#### Protein digestion

Samples were heated for 10 min at 95°C and subjected to 1 min High Intensity Focused Ultrasound (HIFU) at an ultrasonic amplitude of 100 % before centrifugation at 20000 x g for 10 min. The protein concentration was determined using the Lunatic UV/Vis polychromatic spectrophotometer (Unchained Labs) with a 1.10 dilution for each sample. For each sample a volume according to 50 µg of protein was taken and reduced with 5 mM TCEP (tris(2-carboxyethyl)phosphine) and alkylated with 15 mM chloroacetamide at 30°C for 30 min in the dark. Protein clean-up and digestion were done using the single□pot solid□phase enhanced sample preparation (SP3) method using a KingFisher Flex System (Thermo Fisher Scientific) and Carboxylate-Modified Magnetic Particles (GE Life Sciences; GE65152105050250, GE45152105050250) [176, 177]. Samples were diluted with 100% ethanol to a final concentration of 60% ethanol and the following steps were carried out on the robot: collection of beads from the last wash, protein binding to beads, washing of beads in wash solutions 1-3 (80% ethanol) and peptide elution from the magnetic beads using MilliQ water after collecting the digest solution the next day. Protein digestion with a trypsin : protein ratio of 1:50 in 50 mM Triethylammoniumbicarbonat (TEAB) was done offline with a sealed plate on a Thermomixer at 37°C overnight. The digest solution and water elution were combined and dried to completeness.

#### Phosphopeptide enrichment

The phosphopeptide enrichment was performed using a KingFisher Flex System (Thermo Fisher Scientific) and Ti-IMAC HP MagBeads (ReSyn Biosciences). Beads were conditioned following the manufacturer’s instructions, consisting of 3 washes with 400 µl of binding buffer (0.1 M glycolic acid, 80% acetonitrile, 5% TFA). Each sample was dissolved in 150 µl binding buffer and an aliquot of 1 ug per sample was taken for the whole proteome analysis. Beads, wash solutions and samples were loaded into 96 well microplates and transferred to the KingFisher. Phosphopeptide enrichment was carried out using the following steps: binding of the phosphopeptides to the beads (30 min), washing the beads in wash 1, 2 and 3 (wash buffer 1: 0.1 M glycolic acid, 80% acetonitrile, 5% TFA, wash buffer 2: 80% acetonitrile, 1% TFA, wash buffer 3: 10% acetonitrile, 0.2% TFA, 3 min each) and eluting peptides from the beads (80 µl 1% NH4OH in water, 10 min) [177]. To each elution 10 µl of 20% formic acid were added. Phospho-enriched as well as a portion of the small input fraction for full proteome analysis corresponding to 300 ng of peptides were loaded onto Evotips according to the manufacturer’s instructions.

#### LC-MS/MS data acquisition

MS analyses were performed on a timsTOF HT (Bruker) coupled to an Evosep One (EvoSep Biosystems). Samples were separated with the extended Evosep method ‘‘15 samples/day’’ with the analytical column (PepSep, ReproSil C18 3m 120 Å 15cm ID 100 µm) at 50°C. For the dual timsTOF, MS spectra were scanned from m/z 100 to m/z 1700 in ddaPASEF mode (data dependent acquisition Parallel Accumulation Serial Fragmentation). For the ion mobility settings, the inversed mobilities from 1/K0 0.70 Vs/cm2 to 1.50 Vs/cm2 were analyzed with ion accumulation and ramp time of 100 ms, respectively. 1 survey TIMS-MS scan was followed by 10 PASEF ramps for MS/MS acquisition, resulting in a 1.17 s cycle time. Singly charged ions were excluded using the polygon filter mask while active exclusion was set at 0.4 min for ions up to charge state 5. Isolation windows for MS/MS were set to 2.0 Th for precursor ions below and at m/z□700 and 3.0 Th for ions at or above 800 m/z. Collision energy settings were tuned using 20, 25, 54 and 60 eV at 0.6, 0.85, 1.17 and 1.5 1/K0 values. The mass spectrometry proteomics data were handled using the local laboratory information management system (LIMS) [178].

#### Protein and Phosphosite Identification

The acquired shotgun MS data were processed for identification and quantification using Fragpipe 23.1 (MSFragger 4.3, Philosopher 5.1.2, IonQuant 1.11.11). Spectra were searched against a concatenated Uniprot Homo sapiens reference proteome (UP000005640, reviewed canonical version from 2025-10-13 concatenated to its reversed decoyed fasta database and common protein contaminants) with carbamidomethylation of cysteine as fixed, and methionine oxidation and phospho STY (enriched samples only) as variable modifications. Error tolerances were kept at +/-20 ppm for MS1 and 20 ppm for MS2. Enzyme specificity was set to trypsin/P allowing a minimal peptide length of six amino acids and a maximum of two missed cleavages. Label-free quantification, match-between-run and MSBooster (1.3.17, DIA-NN models for ion mobility, retention time and spectra) were activated. Post-search site localization scoring was done with PTMProphet (7.3.0). Liquid chromatography-mass spectrometry (LC-MS) data were processed to identify both total protein abundances and site-specific phosphorylation (PTM). For the total proteome analysis, 7,906 proteins were identified (1.01% decoy sequences; 0.19% contaminants). For the phosphoproteome analysis, 42,477 sequences were identified (0.72% decoy sequences; 0% contaminants). In both datasets, decoy (REV/rev) and contaminant (CON/zz) sequences were retained during initial processing to facilitate a robust re-estimation of false discovery proportions in the final results. All matrices were filtered to include only entries with at least one quantified peptide.

#### Data Pre-processing and Normalization

Raw abundances were processed using the R package prolfqua [179]. To address systematic differences in sample concentration and loading, Variance Stabilizing Normalization (VSN) was applied [180]. This procedure stabilized variance across the intensity range and log2 -transformed the empirical abundances to a scale-free format [181]. Quality control and group consistency were assessed via hierarchical clustering based on Minkowski distance and the coefficient of variation (CV) of abundances. Principal Component Analysis (PCA) was employed to visualize sample similarity and identify potential outliers based on global profiles.

#### Missing Value Analysis and Imputation

Data completeness was evaluated using a dichotomous present/absent analysis. The total proteome dataset contained 2,647 proteins (33.08%) with at least one missing value, while the phosphoproteome exhibited a higher degree of sparsity with 40,619 sequences (94.94%) containing missing observations. Non-detections were assumed to arise primarily from abundances falling below the Limit of Detection (LOD). For proteins or sites where a treatment group lacked any observations, the unobserved group mean was imputed using the mean of the 1% smallest group-averages. This approach enabled the statistical inclusion of binary presence/absence patterns across conditions.

#### Differential Expression Analysis

Differential expression (DE) was modeled using a moderated linear model framework. To account for the paired experimental design, normalized abundances were modeled as a function of the treatment group and donor. Protein and site variances were moderated using an empirical Bayes approach [182], shrinking individual variances towards a global prior to increase statistical power.

#### Proteomics analysis

Scores for all genes in each pathway within the MSigDB Hallmark collection were first calculated using Gene Set Variation Analysis (GSVA) [183]. To visualize global trends and sample grouping, these scores were represented in a heatmap with annotations for both experimental condition and donor. Subsequently, to identify significant biological shifts, a statistical analysis was performed using paired t-tests. In this framework, all statistical comparisons were conducted by testing each individual condition exclusively against the ‘Untreated’ reference. This same paired statistical pipeline was applied to a targeted set of key genes identified in our previous transcriptomics analysis to evaluate their protein-level abundance trends. Finally, to account for multiple comparisons, p-values were corrected using the Benjamini-Hochberg False Discovery Rate (FDR) method.

#### Proteomics and phosphoproteomics integration and differential analysis

To distinguish between changes in signaling activity and changes in protein abundance, total proteome and phosphoproteome datasets were integrated following the MSstatsPTM approach [184]. For the 87.34% of identified phosphoproteins (5,048 proteins) also quantified in the total proteome, phosphorylation site statistics were normalized by their corresponding protein-level changes. The protein-level difference was subtracted from the phosphorylation site difference for each contrast. This adjustment, referred to as Differential PTM Usage (DPU), generates normalized statistics. This integrated analysis reveals whether observed phosphorylation shifts are driven by protein expression changes or by genuine site-specific modification usage, providing deeper insights into cellular regulation.

#### Inference of kinase activity

Kinase enrichment analysis was performed using the KEA3 web tool [185], querying kinase–substrate relationships from the Kinase Library resource [186]. Proteins containing differentially upregulated phosphosites, defined by log2FC ≥ 1 and FDR ≤ 0.05, were submitted as the input set. KEA3 assesses enrichment of kinase-associated substrates using Fisher’s exact test against its default background set, with multiple testing correction performed using the Benjamini–Hochberg method. The odds ratio indicates the enrichment of substrates associated with a given kinase among the input proteins relative to their expected occurrence in the background set.

### Similarity Scores

Two data modalities were analyzed: (i) bulk RNA-sequencing profiles derived from an in-vitro model, and (ii) scRNA-seq obtained from solid tumors of cancer patients. The objective was to quantify the similarity between these datasets. This requires addressing discrepancies arising both from differing measurement technologies and from the biological divergence between experimental models and patient-derived material.

#### Bulk RNA-sequencing data

Each bulk sample was represented as a vector *x* ^(bulk)^ ∈ ℝ^*G*^, where *G* denotes the number of measured genes. Vectors were first scaled such that their *l*_1_ -norm equalled 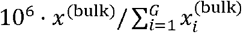. An elementwise log-plus-one transformation was subsequently applied, such that one yields 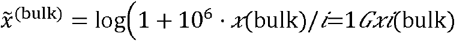.

#### Single-cell RNA-sequencing data

Single-cell profiles – including a clustering thereof – were obtained from the TAM-atlas provided by Coulton et al [21]. The preprocessing applied in the atlas was the following: each cell was scaled to an *l*_1_ norm of 10^6^ followed by an elementwise log-plus-one transformation. The individual datasets constituting the atlas were then harmonized using the RPCA integration workflow implemented in Seurat (v4.2.0). Clustering was performed using the established workflow consisting of the sequential application of the functions RunPCA, FindNeighbors, and FindClusters by the Seurat (v4.2.0) package. For further details on the integration, or clustering workflow, see Coulton et al. [21]. For each cell, the resulting vector representing the normalized, harmonized expression profile will be denoted as 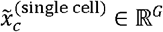, where *c* =1,…, *C* with *C* being the number of cells in the TAM-atlas.

#### Aggregation of Mean Distances

First, we calculated the mean distance between transcriptome profiles for each pair of conditions in the bulk RNA-sequencing data and cell cluster in the TAM atlas. For each pair, we computed the arithmetic mean of the distance values 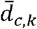, where *c* represents the cluster and *k* represents the condition, resulting in a matrix of mean distances across all cluster-cytokine combinations.

#### Second-Order Difference Analysis

To isolate the effect of specific treatments relative to the baseline, we performed a second-order difference analysis. We defined the untreated condition as the reference state for each cluster. For every condition *k* in cluster *c* apart from the untreated condition *u*, we calculated the difference Δ_*C,k*_ as:

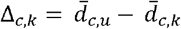

This formulation yields the magnitude of the distance shift relative to the untreated control. Finally, we applied a z-normalization on per condition and afterwards a second per cluster.

#### Statistical Significance via Permutation Testing

To evaluate the significance of the observed distance differences Δ_*c,k*_, we employed a label-shuffling permutation test. For each biological replicate, we randomly reassigned the cytokine condition labels across all cells within a cluster, while strictly preserving the untreated condition as the baseline reference.

This procedure was repeated *N* =1000 to generate a null distribution of distance differences under the hypothesis of no condition effect. For each condition-cluster pair, we calculated an empirical p-values p_*c,k*_ as the fraction of permutations where the absolute value of the permuted difference was greater than or equal to the observed differential distance:

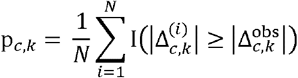

Where I is the indicator function. This approach accounts for donor-specific variance while controlling for random fluctuations in the cluster-cytokine assignments.

## Survival analysis

A pan-cancer survival analysis was performed across 31 TCGA cohorts, non-solid tumor datasets were removed for this analysis. Transcriptomic and clinical data were downloaded from The Cancer Genome Atlas (TCGA) using the TCGAbiolinks v.2.38.0 R package, restricted to Primary Solid Tumor samples. To account for tissue-specific expression variability, thresholds were determined independently for each cancer type. For each gene of interest, patients within each specific cancer cohort were stratified into “High” and “Low” expression groups. The optimal cut-off point to separate high and low expression was calculated using the surv_cutpoint and surv_categorize functions from the survminer v.0.5.1 R package, applying a minimum proportion (minprop) of 0.2 to ensure statistical stability. The analysis was performed for genes *MT1A, MT1DP, MT1E, MT1G, MT1H, MT1JP, MT1L, MT1M, MT1X, MT2A*, and *MTF1*.

Survival probabilities were estimated using the Kaplan-Meier (KM) method using the survival v.3.8.6 R package, with group differences assessed by the log-rank test, the resulting p-values were adjusted for multiple testing using Bonferroni correction. Hazard Ratios (HR) and 95% confidence intervals (CI) were derived from univariable Cox Proportional Hazards regression models.

## Supporting information

Supplementary Data containing Supplementary Tables 1-10

## Data availability

Proteomics and phopshoproteomics data will be made available at the proteomics identifications (PRIDE) database upon publication. Additional analysis results are made available as Supplementary Tables.

## Code availability

Code used in this study is available on the GitHub code repository: <u>GitHub - LuJoHae/Macrophage-Response-To-Tumor-Derived-Cues · GitHub</u>

## END NOTES

## Acknowledgements

The authors gratefully acknowledge the Functional Genomics Center Zurich (FGCZ) of University of Zurich and ETH Zurich, and in particular Dr. Antje Dittmann, Laura Kunz, Dr. Jonas Grossmann and Dr. Witold Wolski for the support on proteomics and phosphoproteomics analyses.

This work was supported by Uniscientia Foundation, Swiss Cancer Foundation, Theron Foundation and Empa-HOCH seed grant.

Figure 1a was created in BioRender. Hast, K. (2026) https://BioRender.com/gwy8bu7

## Author contributions

K. S. led the study; designed and performed experiments; prepared samples for transcriptomic, proteomic, and phosphoproteomic analyses; processed tumor tissues and generated tumor-conditioned media; coordinated functional validation by FACS and cytokine measurements; performed ORA, GSEA, KEA, and TF analyses; visualized the data and prepared the figures; interpreted the results; and wrote the manuscript with input from all authors. L. J. H. performed data integration with the scRNA-seq TAM atlas, conceptualized the methodological approach for dataset comparison, wrote the code for similarity score calculation, analyzed expression levels of individual markers of interest, performed data visualization and statistical analysis, and contributed to writing. A. A.-T. contributed to proteomics data analysis, survival analysis, and writing. R. R.-P. performed transcriptomics data processing, differential expression analysis, TFBS analysis, and contributed to writing. J. H. prepared samples for transcriptomic and proteomic measurements and assisted with macrophage polarization experiments. T. T. and J. B. assisted with proteomics data analysis and interpretation of results. A. P. provided clinical samples, jointly performed preprocessing of tumor tissues, and assisted with the preparation of tumor-conditioned media. E. K. assisted with scRNA-seq data analysis and interpretation of results. V. A.-N., M. R., and K. M.-W. contributed to experimental design, troubleshooting, interpretation of results, and writing. S. T. assisted with interpretation of results, writing, and clinical and biological contextualization. M. N. contributed to experimental design, troubleshooting, interpretation of results, provision of clinical samples, and clinical contextualization. B. S. assisted with experimental design, troubleshooting, and interpretation of results. M. B. conceptualized and supervised the study, assisted with experimental design and interpretation of results, and wrote the manuscript with input from all authors.

